# Binding of the SARS-CoV-2 Spike Protein to Glycans

**DOI:** 10.1101/2020.05.17.100537

**Authors:** Wei Hao, Bo Ma, Ziheng Li, Xiaoyu Wang, Xiaopan Gao, Yaohao Li, Bo Qin, Shiying Shang, Sheng Cui, Zhongping Tan

## Abstract

The pandemic of SARS-CoV-2 has caused a high number of deaths in the world. To combat it, it is necessary to develop a better understanding of how the virus infects host cells. Infection normally starts with the attachment of the virus to cell-surface glycans like heparan sulfate (HS) and sialic acid-containing oligosaccharides. In this study, we examined and compared the binding of the subunits and spike (S) proteins of SARS-CoV-2 and SARS-CoV, MERS-CoV to these glycans. Our results revealed that the S proteins and subunits can bind to HS in a sulfation-dependent manner, the length of HS is not a critical factor for the binding, and no binding with sialic acid residues was detected. Overall, this work suggests that HS binding may be a general mechanism for the attachment of these coronaviruses to host cells, and supports the potential importance of HS in infection and in the development of antiviral agents against these viruses.

## Introduction

The 2019 novel coronavirus (CoV) is the seventh human coronavirus.^1^ It is a deadly virus that is affecting the whole world in an unprecedented way. The global impact of the coronavirus disease 2019 (COVID-19) pandemic is far beyond that of two other major coronavirus outbreaks in the past 20 years, the severe acute respiratory disease (SARS) in 2003^2,3^ and the Middle East respiratory disease (MERS) in 2012.^4^ Given that all three highly pathogenic CoVs were originated from bats and a large number of closely related CoVs are present in bats, future outbreak of this type of zoonotic virus remains possible. In order to avoid facing a similar pandemic in the future, it is necessary to develop a better understanding of these CoVs, especially with regard to effective ways that can help control the current COVID-19 pandemic and prevent the second wave of outbreaks.

Studies showed that the genome of the SARS-CoV-2 has about 80% nucleotide identity with that of SARS-CoV.^5^ The major differences are found in the regions encoding the structural proteins (envelope E, membrane M, nucleocapsid N, and spike S) and accessory proteins (ORF3a/3b, 6, 7a/7b, 8 and 10), not the nonstructural proteins (nsp1 to nsp16). Based on this genetic similarity, the 2019 novel CoV was named by the International Committee on Taxonomy of Viruses (ICTV) as the severe acute respiratory syndrome coronavirus 2 (SARS-CoV-2).^6^ SARS-CoV-2 is more genetically distant from MERS-CoV and shares only about 57% genome homology with MERS-CoV.^5,7^ These similarities and differences are reflected in the viral binding receptors to cell surface receptors. SARS-CoV-2 and SARS-CoV use the same receptor angiotensin-converting enzyme 2 (ACE2) for entry into target cells,^8–10^ whereas MERS-CoV uses dipeptidyl peptidase 4 (DPP4, also known as CD26) as the primary receptor.^11^ However, there are many important findings that cannot be explained by their genetic relations. For example, it was found that SARS-CoV-2 has a relatively lower mortality rate (2%) than SARS-CoV (15%) and MERS-CoV (35%), but is more transmissible among humans.^12–14^ These findings suggest that more investigations other than genetic analysis need to be carried out to improve the understanding of SARS-CoV-2 infection.

In order to efficiently infect host cells, SARS-CoV-2 must bind with cell surface molecules in the lungs and other organs to mediate viral attachment and entry into host cells. Previous studies of many other viruses suggested that SARS-CoV-2 S protein may use other molecules on host cell surface as attachment factors to facilitate binding to the high-affinity receptor ACE2.^15,16^ Examples of such molecules include glycosaminoglycans (GAGs) and sialic acid-containing oligosaccharides.^17–20^ GAGs are primarily localized at the outer surface of cells. Such a location makes them particularly suitable for acting as attachment factors to recruit viruses to cell surfaces.^21,22^ HS is one of the most prevalent types of GAGs in mammals. It is a linear and sulfated polysaccharide that is abundantly expressed on the surface of almost all cell types and in the extracellular matrix.^23,24^ The HS chains are mostly covalently linked as side chains to core proteins to form HS proteoglycans (HSPGs) (Fig. 1).

**Figure 1.**
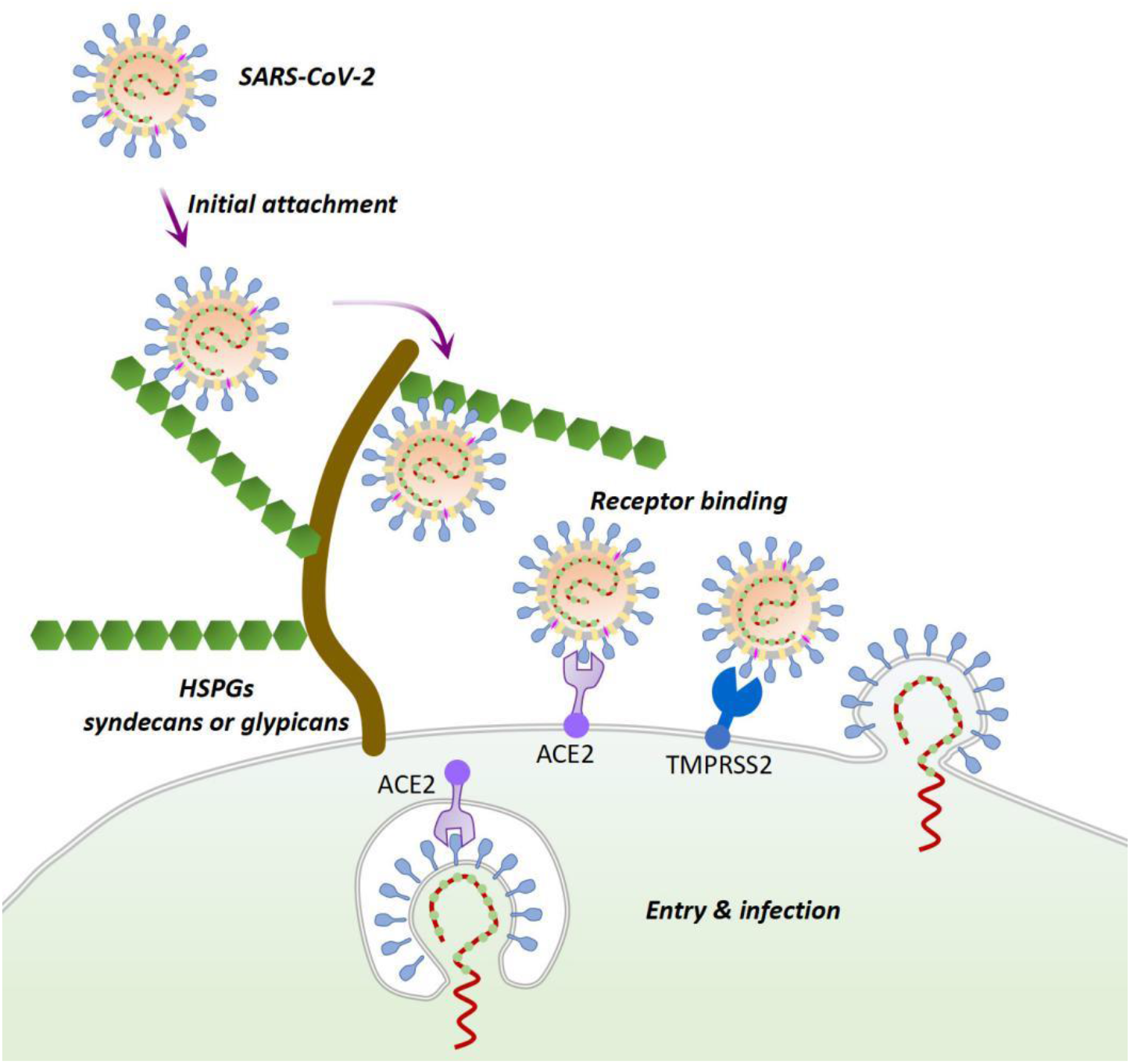
A possible mechanism for SARS-CoV-2 entry and infection. At the early stage of the infection process, SARS-CoV-2 may first interact with the HSPGs on the surface of susceptible cells using the S protein protruding from the virus particle. This initial attachment may promote the subsequent binding of the virus to the high-affinity entry receptor ACE2. The serine protease TMPRSS2 on host cell surface and other host cell proteases may assist in viral entry by cleaving the S protein at the S1/S2 and/or at the S2’ sites.

HS is synthesized in the Golgi apparatus by many different enzymes. During and after its assembly, HS undergoes extensive series of modifications including sulfation, acetylation and epimerization, which leads to glycan structures with high heterogeneity in length, sulfation, and glucuronate/iduronate ratio.^25^ Considerable variation in the sulfation pattern and degree of HS was noted in different species, organs, tissues, and even at different ages and disease stages.^23,24,26^ The sequence and sulfation pattern of HS has been shown to be able to regulate the binding of many viruses to host cells during infection.^27,28^ A similar trend was also observed for sialylation patterns of cell surface oligosaccharides.^17,29,30^ These findings implicate that a possible relationship may exist between the distribution of different types of HS/sialylated glycans and the viral tropism.^31–33^ A better understanding of their relationship can potentially contribute to the design of new antiviral strategies. Although currently the data on the viral tropism of SARS-CoV-2 are limited, results from recent studies suggested that its tropism may not be correlated with the ACE2 expression.^34,35^ Some other factors, such as proteases and glycans, may be the determinant of cellular susceptibility to the infection with this virus.^36^ A recent study suggested that HS may bind to the receptor binding domain (RBD, the C-terminal region of the S1 subunit, Fig. 2) of the SARS-CoV-2 spike protein and change its conformation.^37^ The intriguing possibility that variation in HS and sialic acid characteristics could impact the tropism of viruses prompts us to investigate the binding of SARS-CoV-2 toward a series of HS and sialic acid containing oligosaccharides.^38^

**Figure 2.**
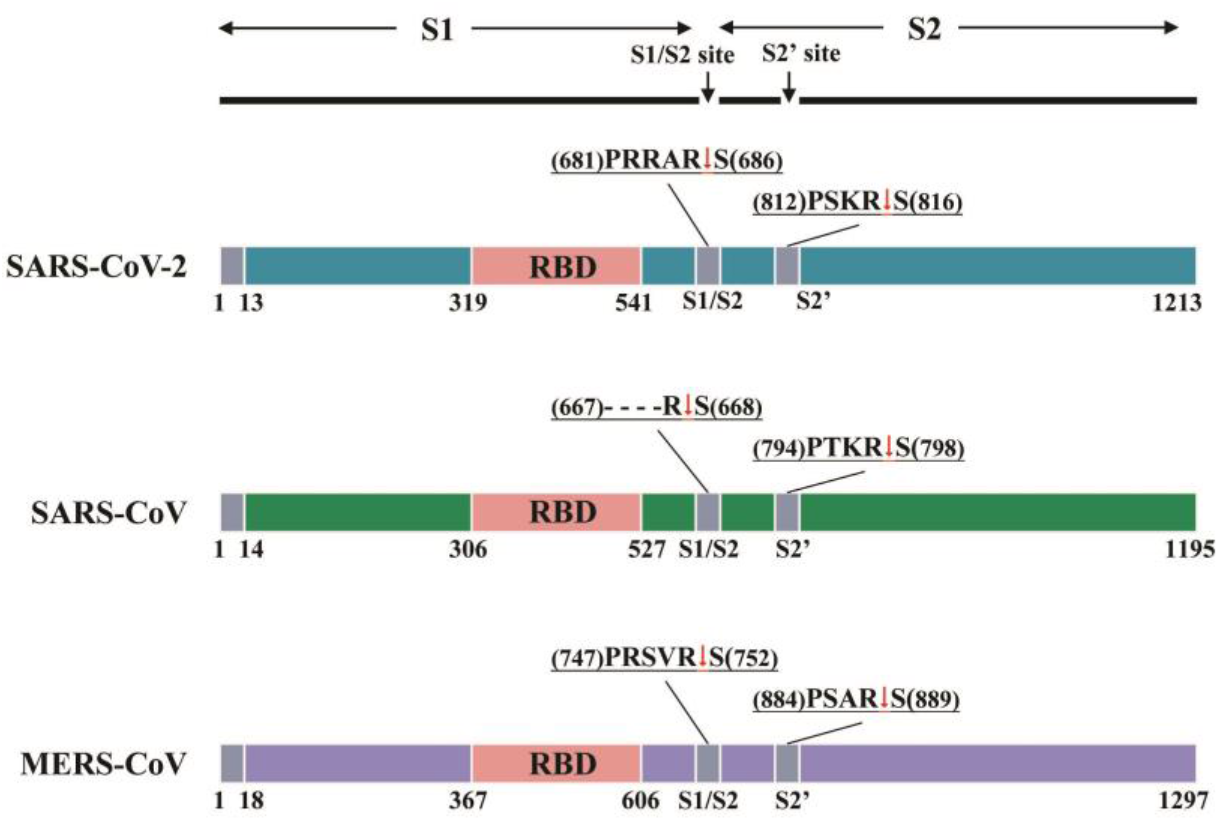
Schematic diagram showing the domain organization of the S proteins of SARS-CoV-2, SARS-CoV and MERS-CoV. The RBD is shown in pink color. The sequences of the S1/S2 and S2’ cleavage sites are given on top of each diagram. The red arrows indicate the cleave points.

In this study, we systematically examined and compared the binding of the SARS-CoV-2 S protein subunits, full-length molecule and its trimer to different HS using microarray experiments (Fig. 2). Our results suggested that all the tested protein molecules were capable of binding HS oligomers and had similar binding preferences, with higher affinity toward HS forms with higher degrees of sulfation. The binding was also shown to be positively related to the level of 6-O-sulfation. The HS chain length seemed to be not critical. A long heparin molecule (a highly sulfated HS) and a synthetic heparin pentasaccharide (Fondaparinux sodium) were demonstrated to have similar binding affinity to these protein molecules. Moreover, our study suggested that the SARS-CoV-2 S protein might not be able to bind sialic acid residues, or the binding might be too weak to be detected by the microarray technology, and the S proteins of SARS-CoV and MERS-CoV have similar HS and sialic acid binding properties as those of the SARS-CoV-2 S protein. Overall, our study laid a foundation for future studies to explore whether the binding specificity to HS can serve as an important contributor to the viral tropism of SARS-CoV-2 and to explore the possibility of exploiting HS for therapeutic strategies.

## Results

### Binding of the subunits and S proteins of SARS-CoV-2, SARS-CoV, and MERS-CoV to a HS microarray

An early study has suggested that the RBD of SARS-CoV-2 S protein might bind heparin.^9^ In order to determine if there is any preference of the SARS-CoV-2 S protein for particular HS structures, we first investigated the binding of the RBD (Here termed as SARS-CoV-2-RBD) to a HS microarray containing 24 synthetic heparan sulfate oligosaccharides. These oligosaccharides have systematic differences in their length, monosaccharide composition, and sulfation pattern (Fig. 3). The microarray experiment was performed using a previously established standard protocol.^29,39,40^ Briefly, the proteins were labeled with Cy3 fluorescent dye and incubated with the microarray at different concentrations. After washing away the unbound HS molecules, a highly sensitive fluorescence method was used to detect the binding of the SARS-CoV-2-RBD to HS. Under the experimental conditions, the binding can be detected at concentrations higher than 0.5 μg/ml. Increasing the concentration of the protein was not found to produce noticeable changes in binding (Supporting Information, Fig. S1).

**Figure 3:**
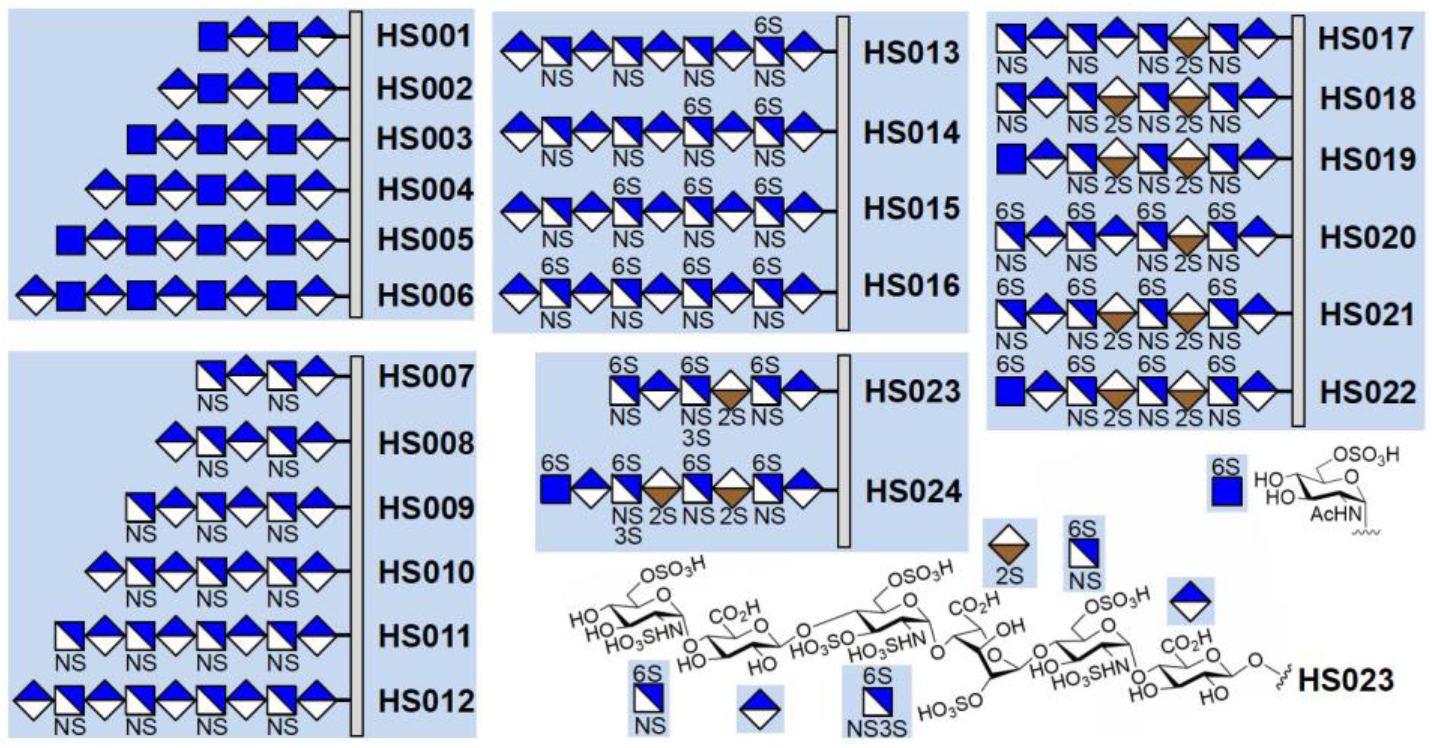
The numbering and structures of the HS oligosaccharides on the microarray. Each HS chain is covalently attached to the microarray slide via the reducing end. The HS oligosaccharides can be roughly divided into five groups according to their differences in the monosaccharide compositions and sulfation patterns.

Quantification of fluorescence revealed that the SARS-CoV-2-RBD is able to bind to almost half of the molecules on the microarray, and not surprisingly, the binding is strongly affected by the sulfation level, which is a trend that has been previously noted for many HS-binding proteins.^41–43^ As shown in Figure 4a, the HS oligosaccharides with higher sulfation degree, **HS020-HS024** (the number of sulfate groups per monosaccharide unit >1.00), exhibit higher fluorescence intensity. The highest fluorescence intensity is observed for **HS023** (1.35 sulfate groups per monosaccharide), which is followed by those of **HS021** and **HS024** (1.25 sulfate groups per monosaccharide) and those of **HS020** and **HS022** (1.15 sulfate groups per monosaccharide). Binding of SARS-CoV-2-RBD to HS seems to not be affected by the monosaccharide composition (compare **HS020** with **HS022**, and **HS021** with **HS024**).

**Figure 4.**
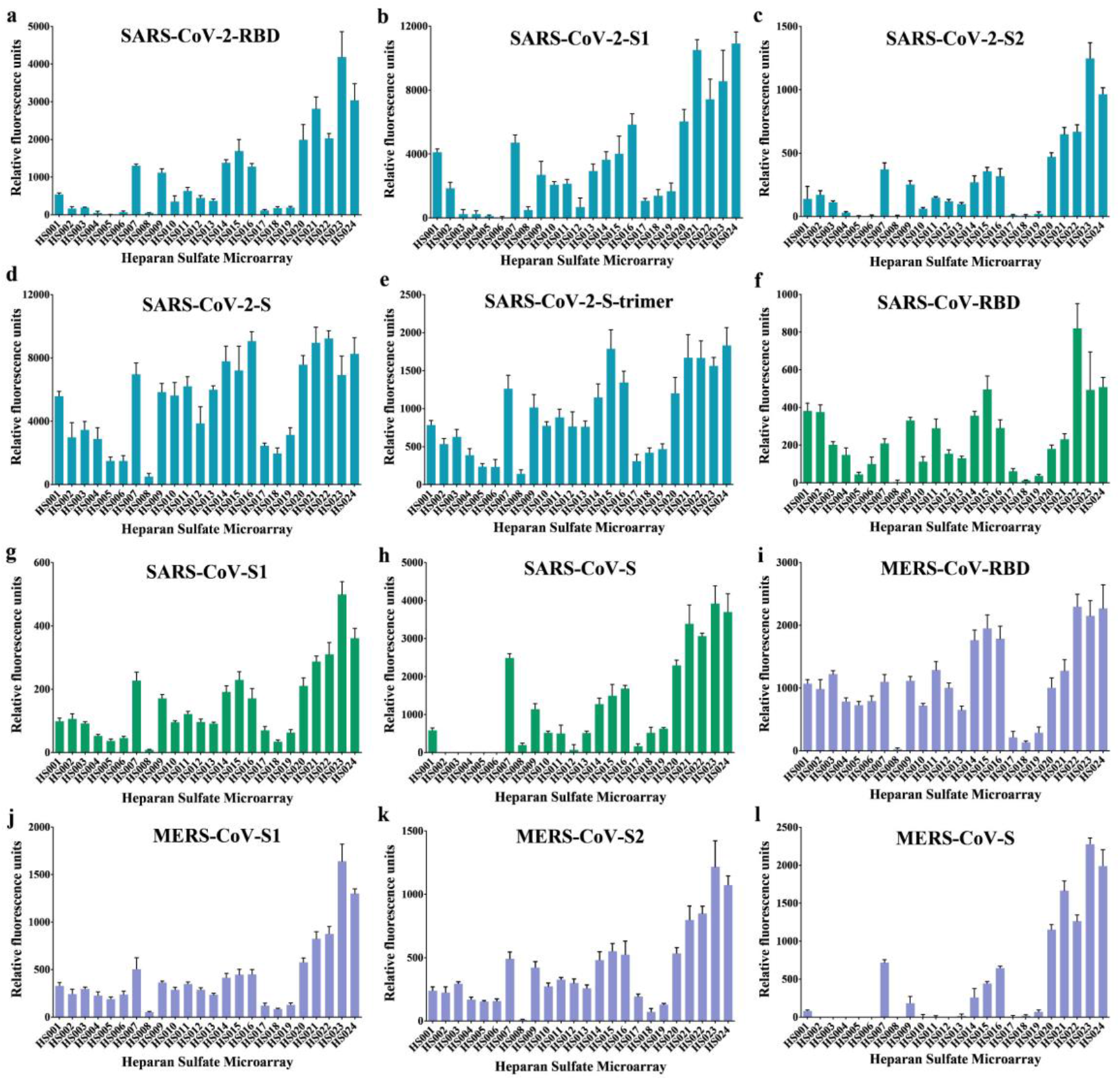
The contributions of different HS characteristics to the binding to (a) SARS-CoV-2-RBD at the concentration of 2.5 μg/ml, (b) SARS-CoV-2-S1 at the concentration of 2 μg/ml, (c) SARS-CoV-2-S2 at the concentration of 2 μg/ml, (d) SARS-CoV-2-S at the concentration of 2 μg/ml, (e) SARS-CoV-2-trimer at the concentration of 2 μg/ml, (f) SARS-CoV-RBD at the concentration of 2 μg/ml, (g) SARS-CoV-S1 at the concentration of 2 μg/ml, (h) SARS-CoV-2-S at the concentration of 2 μg/ml, (i) MERS-CoV-RBD at the concentration of 4 μg/ml, (j) MERS-CoV-S1 at the concentration of 4 μg/ml, (k) MERS-CoV-S2 at the concentration of 2 μg/ml, (l) MERS-CoV-S at the concentration of 2 μg/ml. The fluorescence intensity was measured at an excitation wavelength of 532 nm. All error bars are standard deviation of more than three replicates.

The oligosaccharides with relatively lower sulfation levels (**HS001-HS019**, the number of sulfate groups per monosaccharide unit <1.00) have lower or almost no fluorescence signals. An analysis of the effect of the variation in sulfation revealed that the position of sulfation is another factor that strongly influence the binding. As shown in the Figure 4, removal of the 6-O-sulfate group from the glucosamine units significantly reduced the binding (compare **HS017** with **HS020, HS018** with **HS021,** and **HS019** with **HS022**), suggesting that the 6-O-sulfate in HS plays a crucial role in determining its interaction with SARS-CoV-2-RBD. The importance of the 6-O-sulfate for binding is further supported by comparing the binding of **HS012, HS013, HS014, HS015,** and **HS016**, which shows that the one-by-one addition of sulfate to the 6-O-position of the glucosamine residues gradually increased the binding of HS with SARS-CoV-2-RBD. The microarray study also indicates that the binding is not positively related to the length of the HS chains. Short HS oligosaccharides could have comparable or even better binding properties (compare **HS001-HS006, HS007-HS0012**, and **HS020-HS024**).

The binding results of the SARS-CoV-2-S1 follow a similar trend as those of the SARS-CoV-2-RBD, with only minor differences for a few HS molecules like **HS001, HS007, HS016, HS021** and **HS024** (Fig. 4b). This is consistent with the assumption that RBD is the major determinant for viral S protein binding to HS.

Examination of the sequence of the SARS-CoV-2 S protein revealed the presence of a potential cleavage site for furin proteases (RRARS) at the S1/S2 boundary.^9^ This is similar to the S protein of MERS-CoV, which also contain a furin consensus sequence RXXR (Fig. 2). Furin cleavage can occur in the secretory pathway of infected cells and breaks the covalent linkage between the S1 and S2 subunits.^44^ The S2 subunit functions to fuse the virus to the host cell. It may interact with HS on the cell surface to facilitate the fusion process. Therefore, to determine the role of HS in SARS-CoV-2 infection, it is also important to investigate the binding of the S2 subunit to HS. Very interestingly, our results indicate that the SARS-CoV-2-S2 can also bind to HS, with the binding preferences similar to those of the SARS-CoV-2-RBD and the SARS-CoV-2-S1 (Fig. 4c). This suggests that HS may also play an important role during viral membrane fusion after the S1 subunit is removed from the S protein.

We also further investigated the binding full-length S protein and its trimer to HS using the same microarray. Our data showed that, although the binding to **HS020-HS024** still remains the highest, the full-length S protein has increased binding to HS with slightly less sulfation, particularly the molecules in groups **HS007-HS012** and **HS013-HS016** (Fig. 4d, 4e). Additionally, we found that the binding preferences of the RBDs, S1 subunits and full-length S proteins of SARS-CoV and MERS-CoV are similar to those of the subunits and S protein of SARS-CoV-2, although small differences are observed (Fig. 4f-k). Because the MERS-CoV S protein also contains a furin cleavage site, we studied the binding of its S2 subunit to the HS microarray. Once again, the results are similar to those of the S2 subunit of SARS-CoV-2. Overall, these data support the involvement of HS in the binding with the subunits and S proteins of SARS-CoV-2, SARS-CoV and MERS-CoV and support the importance of HS for the infection of these coronaviruses.

### Measurement of the dissociation constants of a long heparin molecule and fondaparinux to the immobilized subunits and S proteins of SARS-CoV-2, SARS-CoV, and MERS-CoV by surface plasmon resonance (SPR)

To determine the relative binding strength of the subunits and S proteins of SARS-CoV-2, SARS-CoV, and MERS-CoV, we measured and compared the dissociation constants (K_D_) of the protein molecules studied in the microarray experiment using a real-time SPR-based binding assay. A commercially available porcine heparin from Sigma-Aldrich was used as the first binding partner for the measurement. It is a mixture of highly sulfated HS, with most chains in the molecular weight range of 17 to 19 kilodalton. This long heparin molecule is more similar to the HS molecules on the host cell surface than those on the microarray. In the SPR assay, protein molecules were covalently linked to the surface of the CM5 (carboxymethylated dextran) sensor chips by amine coupling. The heparin in various concentrations was then flowed over the immobilized proteins. The changes in refractive index by molecular interactions at the sensor surface were monitored and the dissociation constants were obtained by fitting the results using the software available in the SPR instrument (Fig. 5).

**Figure 5.**
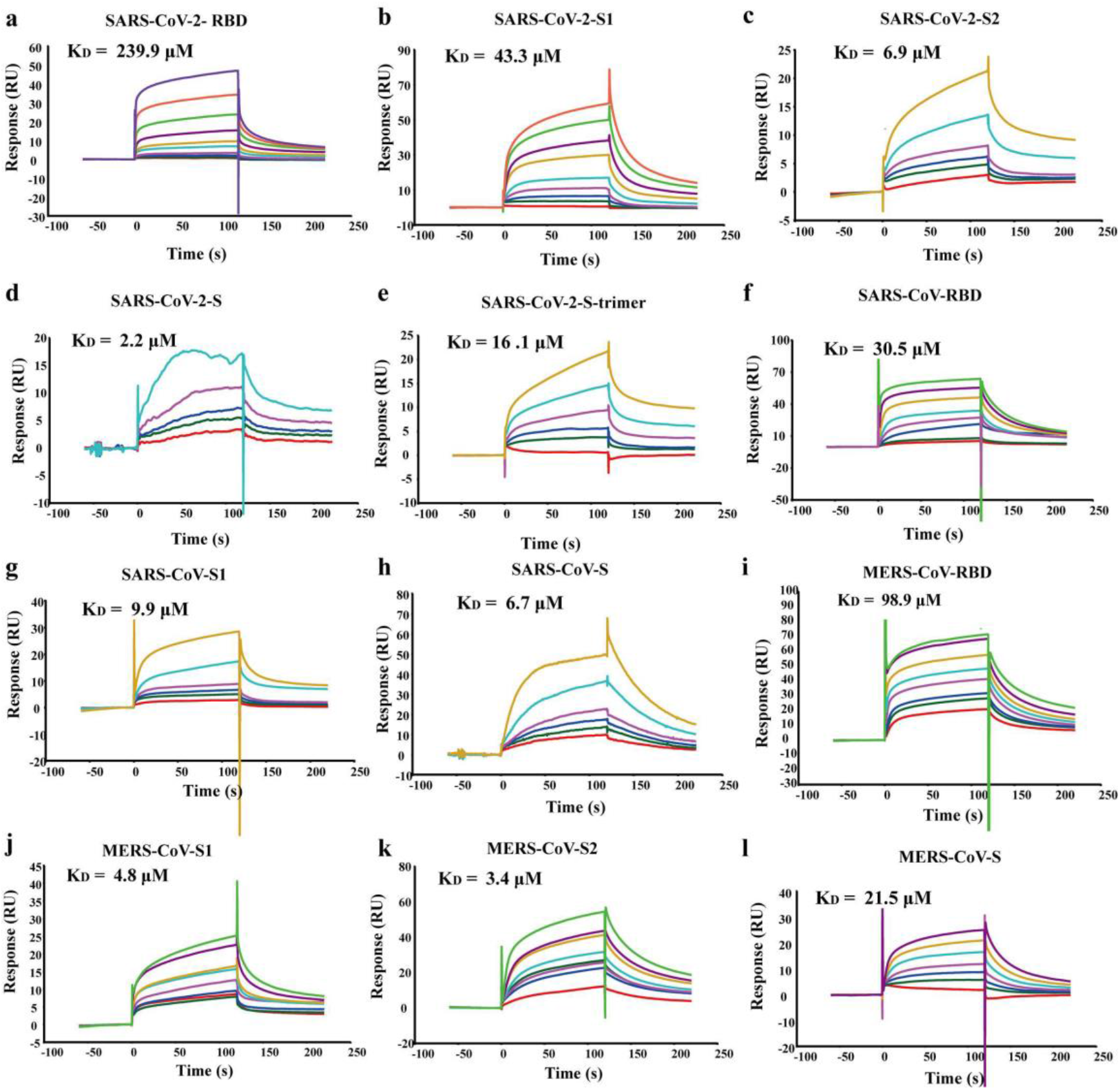
SPR sensorgrams showing the interactions between the long heparin molecule and immobilized proteins. The name and K_D_ value of each protein are presented on top of the corresponding sensorgrams.

The comparison of the dissociation constants revealed that all the tested protein molecules can bind to the heparin, but their binding affinities are relatively low (K_D_s are at the micromolar level). This agrees well with previous observations that HS is a weak binder to viral S proteins.^45^ The results also showed that RBD has the lowest binding affinity among the tested protein molecules of SARS-CoV-2 (Fig, 5a). The S protein trimer and S1 subunit have relatively lower binding affinity in comparison with the fulllength S protein and S2 subunit, respectively (Fig, 5b-e). Very similar trends were also observed for differences in the binding affinities of the tested protein molecules of SARS-CoV and MERS-CoV (Fig, 5f-l).

In addition to the long heparin molecule, we also measured the binding of the protein molecules to fondaparinux (Arixtra®), which is an ultralow-molecular-weight-heparin (ULMWH) containing five monosaccharide units (Fig. 6). It has a well-defined chemical structure and is currently the only ULMWH that has been clinically approved as an anticoagulant. Fondaparinux is very similar to **HS023** in size, monosaccharide composition, and sulfation position and degree. The results showed that the K_D_ values of fondaparinux binding to the tested protein molecules are in the same range as those for the long heparin. This agrees well with the observation in the microarray study, suggesting that the length of the HS chains is not a determining factor for binding. Despite the similarity, there is a subtle but noticeable difference, which is that the binding affinities of the S1 subunit and RBD to fondaparinux are quite similar.

**Figure 6.**
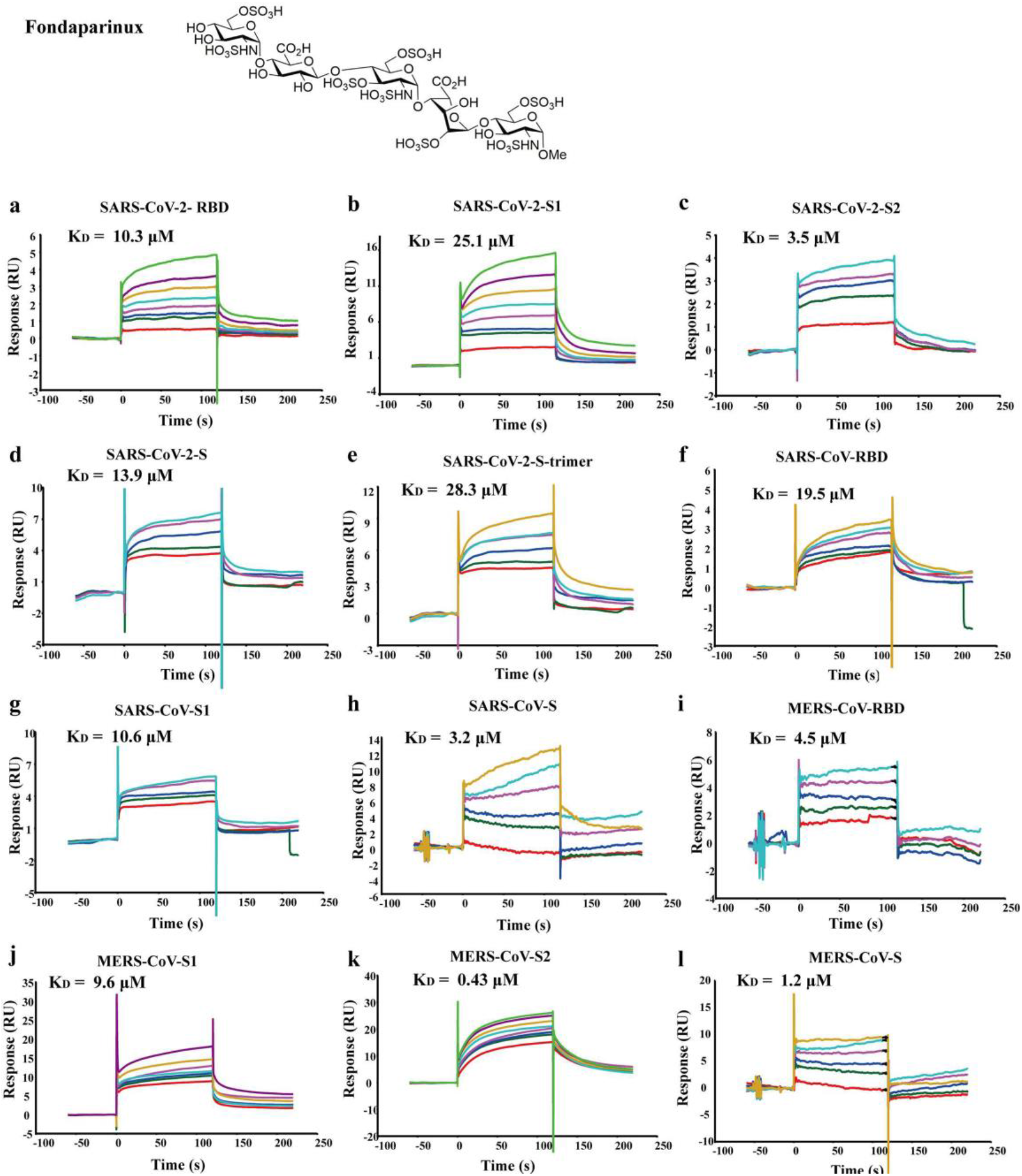
The structure of fondaparinux and the SPR sensorgrams showing the interactions between fondaparinux and immobilized proteins. The name and K_D_ value of each protein are presented on top of the corresponding sensorgrams.

### Binding of the RBD, S1 subunit and S protein of SARS-CoV-2, and S proteins of SARS-CoV and MERS-CoV to sialylated glycan microarrays

Many human viruses can interact with sialic acid-containing glycans present on the cell surface.^17,20^ Such an interaction is normally mediated by the N-terminal domain of the S1 subunit.^46^ In order to find out if SARS-CoV-2 can bind to sialic acid residues, we carried out microarray analyses of its S protein and subunits. The first microarray used contains 100 different N-glycans that may be found on the surface of cells. 49 of them are terminated with α2,3- and α2,6-linked sialic acid, also known as N-acetylneuraminic acid (Neu5Ac), 8 with α2,3- and α2,6-linked N-Glycolylneuraminic acid (Neu5Gc), and the rest with other glycan residues (Supporting Information, Table S1). The experiment was performed in the similar way as described for the HS microarray study. The microarray results showed that both the SARS-CoV-2-RBD and SARS-CoV-2-S1 gave no binding signal, suggesting that they may not be able to interact with sialylated N-glycans or the binding signal is too low to be detected. In order to confirm this finding, we also investigated the binding of the full-length S proteins of SARS-CoV-2, SARS-CoV and MERS-CoV to more sialylated glycans, including sialylated N- and O-linked glycans and glycolipid glycans (Supporting Information, Tables S2-4), but again no specific binding was detected.

## Discussion

For a virus like SARS-CoV-2 to establish infection, it must first attach itself to the surface of target cells in different organs and tissues. The S protein plays an essential role in this attachment process. Recently, the structure of SARS-CoV-2 S protein in the prefusion conformation was determined by the cryo-EM technique.^10^ It shows that the overall structure of the SARS-CoV-2 S protein is very similar to that of the closely related SARS-CoV S protein, which is organized as a homotrimer. Each monomer can be divided into an N-terminal receptor-binding S1 subunit and a C-terminal fusion-mediating S2 subunit. The S1 subunits are located at the apex of the spike, making them more accessible for binding to the proteinaceous receptor, ACE2. Although similar, there are some notable differences between the SARS-CoV-2 and SARS-CoV S proteins.^8,10^ For example, the key amino acid residues involved in the binding of the SARS-CoV-2 S protein to ACE2 are largely different from those of SARS-CoV.^9,47,48^ These differences may be related to the observed higher binding affinity of the SARS-CoV-2 S protein to ACE2.

Another important difference between the SARS-CoV-2 and SARS-CoV S proteins is that the former protein contains a multibasic protease recognition motif (RRARS) at the junction of S1 and S2.^9^ A multibasic cleavage site (RSVRS) was also identified in the MERS-CoV S protein (Fig. 2).^49^ The SARS-CoV S protein only has a monobasic amino acid (SSLRS) at the same site. The multibasic site can be processed by furin or related proprotein convertases, which are widely expressed in different tissues, before the virus is released from the host cell. By contrast, the monobasic site can be cleaved by TMPRSS2 or other cell-surface proteases (whose expression is confined to certain tissues) only after the virus is released from the host cell. It was reported that the cleavage at the junction of S1 and S2 activates the S protein for virus-cell fusion. Thus, the presence of the multibasic cleavage site may partially account for the enhanced infectivity and tropism of SARS-CoV-2 relative to SARS-CoV for human cells.^10,50^ However, what is vague is how the virus attaches to the host cells after losing the S1 subunit, which is responsible for the binding to the proteinaceous receptor.

In addition to binding protein-based receptors, many viruses can interact with cell surface glycans, including GAGs and sialic acid-containing oligosaccharides. Depending on the virus, the glycan molecules can act as attachment factors, coreceptors or primary receptors.^51^ Viruses typically bind GAGs through non-specific charge-based interactions. As one of the most abundant GAGs, HS appears to be the preferred binding partner for many viruses.^33,52–55^ Sialic acids are normally terminal monosaccharide residues linked to glycans decorating cell surface glycoproteins, glycolipids, or other glycoconjugates.^20,56^ In general, the interactions of viruses with HS or sialic acids are responsible for the first contact with host cells. Such contact may serve to concentrate viruses on the surface of target cells, facilitate their binding to more specific high-affinity protein receptors and/or promote their entry into host cells.^57,58^ It has been demonstrated that virus binding and infection can be reduced by enzymatic removal HS or sialic acid from cell surface, or by treating virus with soluble HS or multivalent sialic acid conjugates.^30,59–61^ Therefore, in order to better understand and treat COVID-19, it is necessary to carry out research to investigate the possible interactions between SARS-CoV-2 and HS and sialic acid-containing glycans in the forms of separate subunits and full-length proteins, and to assess if such interactions could represent a target for therapeutic intervention.

Similar to studies that have been successfully conducted for many other viruses, we used the microarray and SPR technology to study the binding of the RBDs, S1/S2 subunits and full-length S proteins of SARS-CoV-2, SARS-CoV and MERS-CoV, and a trimer of the SARS-CoV-2 S protein to HS and sialic acid.^59^ The microarray results showed that all the tested protein molecules can bind to about half of the oligosaccharides on the HS microarray. In contrast, only background levels of fluorescence were detected on various sialylated glycan microarrays. This observation suggests all the tested protein molecules are able to bind to HS^45,62^ and may not bind to and/or have very low binding affinity to sialylated glycans. This well agrees with previous studies showing that SARS-CoV can bind to HS^62^ and the binding of the MERS-CoV S protein to sialic acid-containing glycans can only be detected after the S1 subunit was attached to a nanoparticle to enhance its avidity via multivalent interactions.^17^

Our results also suggested that the binding of the tested proteins to HS is related to HS sulfation position and degree. It seems that more 6-O-sulfate groups and higher sulfation degree normally lead to better binding. Because HSPGs exhibit different sulfation patterns in different tissues, such a binding specificity may contribute to the tropism of SARS-CoV-2 for human cells.^63^ The length of HS appears not to be a critical factor for the binding. Short HS chains could have comparable binding specificity and signals. The comparison of the SPR K_D_ values obtained for the long heparin molecule and fondaparinux further support this finding. It implies that it may be possible to reduce the attachment of SARS-CoV-2 to the surface of host cells by low-molecular-weight-heparin (LMWH). This is in agreement with a recent study showing that LMWH treatment may be associated with better prognosis in some severe COVID-19 patients.^64^ While these initial findings are encouraging, further research is required to determine if the binding to HS could affect the tropism and pathogenesis of SARS-CoV-2 and to determine if HS could be used for the inhibition of the infection of this virus.

Our SPR data of the tested protein molecules also showed that the S2 subunits could have similar or better binding affinity for HS as compared to those of the S1 subunits. This finding suggests that the cleaved S proteins of SARS-CoV-2 and MERS-CoV may depend on HS for interaction with the host cells during viral membrane fusion.

In parallel with our study, the Linhardt and Boons research teams also conducted studies to investigate the binding of S proteins to HS.^45,65^ The absolute values of the dissociation constants determined in our experiments are largely different from those presented by these two teams. This can be seen from the binding of the full-length S protein of SARS-CoV-2, which was studied by all three teams (Supporting Information, Table S5). Our K_D_ is one order of magnitude higher than that reported by the Boons team, which in turn is three orders of magnitude higher than that reported by the Linhardt team. The differences in results may be due to the method of analysis and/or the experimental materials used in the studies. In our study, the tested proteins were immobilized on the surface of CM5 sensor chips, while in their experiments, biotinylated heparin molecules were immobilized on the chips. At the same time, because the S glycoproteins and the heparin molecules were obtained from different sources, their composition may be different from each other.

In conclusion, through our study, we provided experimental evidence for whether or not the S protein of SARS-CoV-2 can bind to two types of cell-surface glycans, HS and sialic acid-containing glycans, which are commonly utilized by human viruses for attachment to target cells. Our data revealed that the SARS-CoV-2 S protein can weakly bind to HS in a sulfation-dependent manner. No binding with sialic acid residues was detected using the microarray assay. The results suggest that HS may act as an attachment factor to concentrate the virus at the cell surface and affect its tropism. Through comparison, we found that the S proteins of SARS-CoV and M ERS-CoV have similar binding properties to HS as that of the SARS-CoV-2 S protein, indicating that HS binding may be a conserved feature for these three types of coronaviruses. Our data also revealed that the S2 subunits could bind equally well as the S1 subunits to HS. This binding may be an important element for viral attachment to the host cell surface after the removal of the N-terminal receptor-binding domains by protease cleavage. Overall, our findings support the potential importance of HS in SARS-CoV-2 infection and in the development of antiviral agents.

## Methods

### Reagents and cell lines

High Five^™^ Cells for baculovirus expression were purchased from ThermoFisher Scientific and maintained in Express Five^™^ Medium. The recombinant SARS-CoV-2-S1 (16-685), SARS-CoV-2-S (16-1213, R683A, R685A), SARS-CoV-2-S-trimer (16-1213, R683A, R685A), SARS-CoV-RBD (306-527), SARS-CoV-S1 subunit (14-667), SARS-CoV-S (14-1195, R677A) with C-terminal His-tags were purchased from ACROBiosystems. The recombinant SARS-CoV-2-S2 (686-1213), MERS-CoV-RBD (367-606), MERS-CoV-S1 (1-725), MERS-CoV-S2 (726-1296), MERS-CoV-S (1-1297) with C-terminal His-tags were purchased from Sino Biological. The heparin sodium salt from porcine intestinal mucosa was purchased from Sigma-Aldrich/Merck. Fondaparinux (sodium salt) was purchased from Macklin Biochemical Co., Ltd (Shanghai, China). The glycan microarray experiments were performed by Creative Biochip Ltd (Nanjing, China).

### Expression and purification of SARS-CoV-2-RBD

DNA containing the coding sequence for an N-terminal hemo signal peptide, the receptor binding domain (RBD, residues 319-541) of SARS-CoV-2 S protein and a C-terminal polyhistidine tag was amplified and inserted into a pFasebac1 vector for expression in High-5 insect cells using the Bac-to-Bac expression system (Invitrogen). The resulting recombinant protein, termed SARS-CoV-2-RBD-His, was secreted into cell culture medium, and subsequently purified on a nickel-nitrilotriacetic acid (Ni-NTA) affinity column, followed by a Superdex 200 gel filtration column (GE Healthcare). The final buffer for the protein contains 10 mM Hepes (pH=7.0) and 100 mM NaCl. The purified SARS-CoV-2-RBD-His was concentrated to 3.5 mg/ml and flash frozen in liquid N2 and stored at −80 degrees Celsius.

### Binding of recombinant proteins to glycan microarrays

The tested protein molecules were first labeled with Cy3 fluorescent dye (10 mg/ml in DMSO). After dialysis, they were incubated at different concentrations with microarrays for 1 hour in the dark at room temperature. After incubation, the microarray slides were gently washed using washing buffer (20 mM Tris-Cl containing 0.1% tween 20, pH 7.4) to remove unbound proteins. Finally, the slides were scanned with a microarray scanner LuxScan-10K/Aat an excitation wavelength of 532 nm and evaluated by the Microarray Image Analyzer software.

### Measurement of dissociation constants by surface plasmon resonance (SPR)

The SPR measurements were performed using BIAcore T200 and S200 (GE healthcare). First, the carboxymethyl dextran matrix on CM5 sensor chip (GE Healthcare) was activated by injection of a 1:1 mixture of 1-ethyl-3-(3-dimethylaminopropyl) carbodiimide (EDC) and N-hydroxysuccinimide (NHS). Recombinant proteins in 10 mM acetate buffer (pH 4.5, GE Healthcare) was then injected over the chip surface at a flow rate of 30 μl/min to couple the amino groups of the recombinant proteins to the carboxymethyl dextran matrix. After the coupling reaction, the remaining activated ester groups were deactivated by ethanolamine. The binding study was carried out at 25°C in PBS-P buffer (GE Healthcare). The heparin molecule or fondaparinux at different concentrations were flowed over the immobilized recombinant proteins at a flow rate of 30 μl/min with a contact time of 60 s and a dissociation time of 100 s. The surface was regenerated by injection of 10 mM Glycine-HCl (pH 2.5) at a flow rate of 30 μl/min for 30 s. Data was collected and analyzed by BIA evaluation software (GE Healthcare).

## Author Contributions

All authors have given approval to the final version of the manuscript. W.H., B.M., S.C., Z.T. designed research; W.H., B.M., Z.L., X.W., X.G., Y.L., B.Q., S.S. performed research; W.H., B.M., S.C., Z.T. analyzed data; W.H., B.M., S.S., S.C., Z.T. wrote the paper.

## Acknowledgement

We would like to thank the National Natural Science Foundation of China (Grant number: 91853120), the National Major Scientific and Technological Special Project of China (Grant number: 2018ZX09711001-013), the National Key R&D Program of China (Grant number: 2018YFE0111400), the State Key Laboratory of Bioactive Substance and Function of Natural Medicines, Institute of Materia Medica, and the Chinese Academy of Medical Sciences and Peking Union Medical College CRP-ICGEB Research Grant 2019 (Grant number: CRP/CHN19-02) for funding. We also thank Guangzhi Shan and Zhiling Zhu for their help in the SPR studies and Kai Zhang and Wenjie Zhou for their support to the research.

---

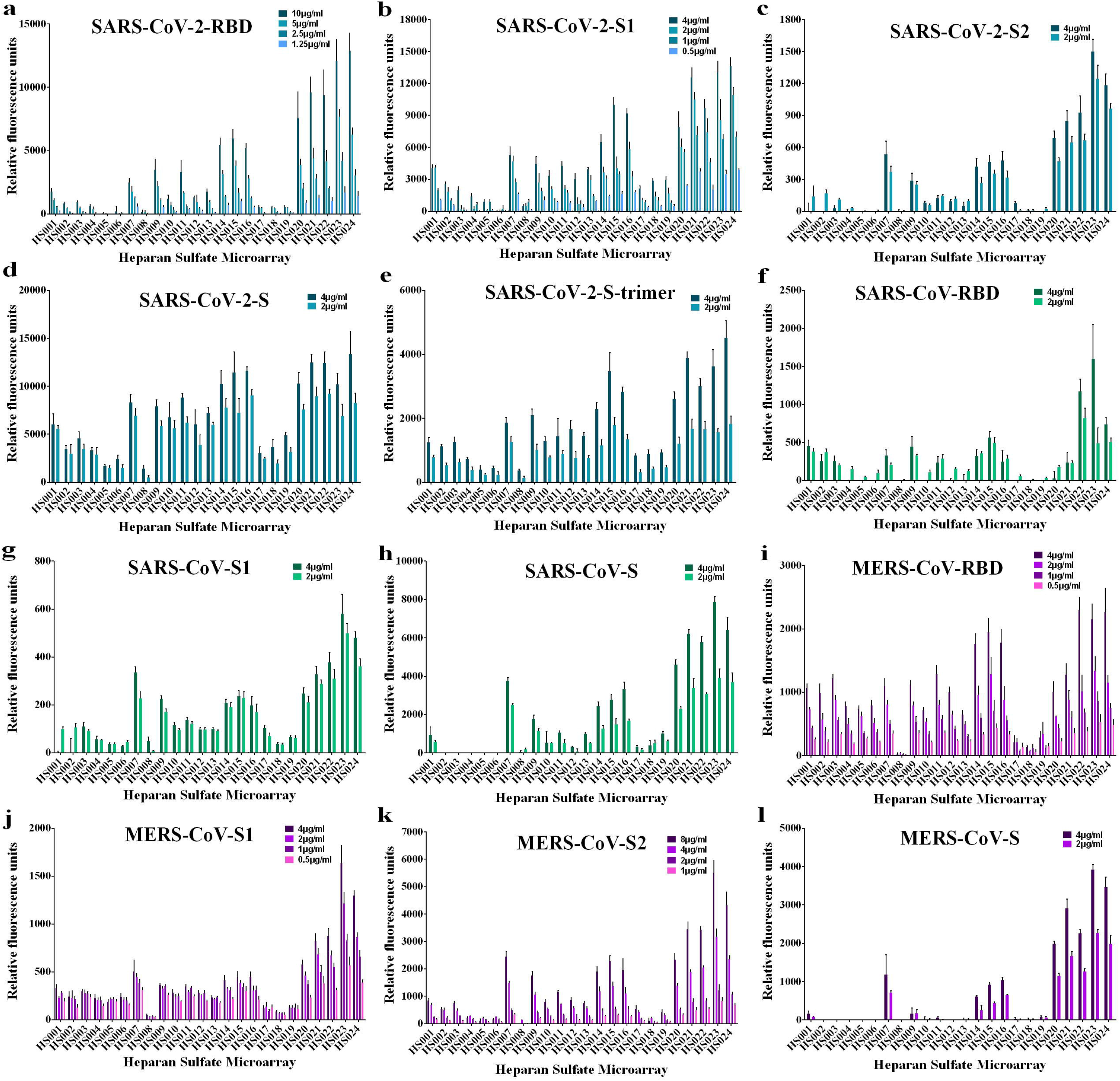

