## supplemental information for "Binding of the SARS-CoV-2 Spike Protein to Glycans"

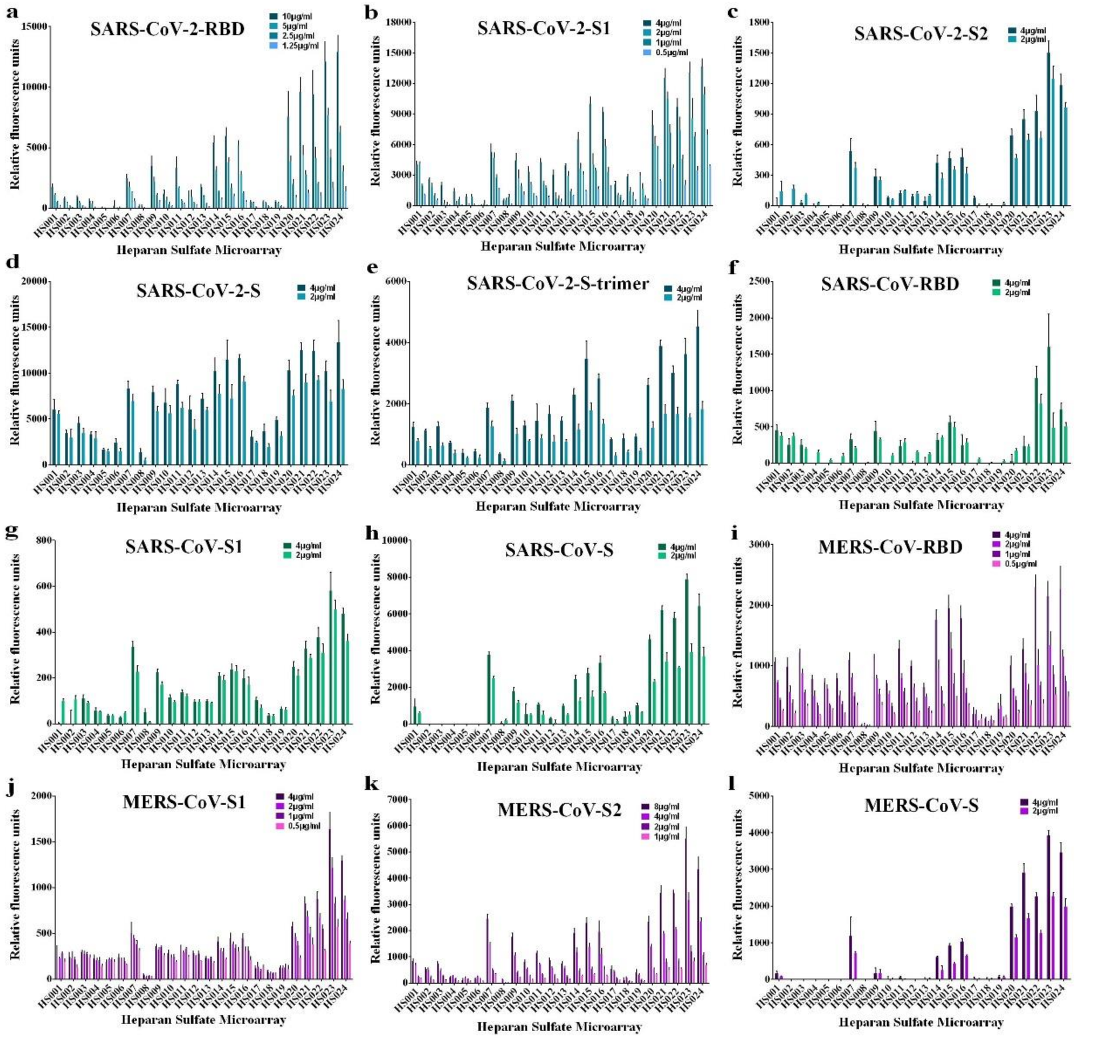


**Figure S1.** Binding of recombinant proteins to the HS microarray at different concentrations. All error bars are standard deviation of more than three replicates.

**Table S1.** The numbering and structures of the N-glycan microarray.

| No. | ID | Structure |
|---|---|---|
| 1 | N000 | GlcNAcβ1-2Manα1-6(GlcNAcβ1-2Manα1-3)Manβ1-4GlcNAcβ1-4GlcNAc- |
| 2 | N001 | Galβ1-4GlcNAcβ1-2Manα1-6(Galβ1-4GlcNAcβ1-2Manα1-3)Manβ1-4GlcNAcβ1-4GlcNAc- |
| 3 | N002 | Neu5Acα2-3Galβ1-4GlcNAcβ1-2Manα1-6(Neu5Acα2-3Galβ1-4GlcNAcβ1-2Manα1-3)Manβ1-4GlcNAcβ1-4GlcNAc- |
| 4 | N003 | Neu5Acα2-6Galβ1-4GlcNAcβ1-2Manα1-6(Neu5Acα2-6Galβ1-4GlcNAcβ1-2Manα1-3)Manβ1-4GlcNAcβ1-4GlcNAc- |
| 5 | N004 | Galβ1-4(Fucα1-3)GlcNAcβ1-2Manα1-6(Galβ1-4(Fucα1-3)GlcNAcβ1-2Manα1-3)Manβ1-4GlcNAcβ1-4GlcNAc- |
| 6 | N005 | Neu5Acα2-3Galβ1-4(Fucα1-3)GlcNAcβ1-2Manα1-6(Neu5Acα2-3Galβ1-4(Fucα1-3)GlcNAcβ1-2Manα1-3)Manβ1-4GlcNAcβ1-4GlcNAc- |
| 7 | N010 | Manα1-6(Manα1-3)Manα1-6(GlcNAcβ1-2Manα1-3)Manβ1-4GlcNAcβ1-4GlcNAc- |
| 8 | N011 | Manα1-6(Manα1-3)Manα1-6(Galβ1-4GlcNAcβ1-2Manα1-3)Manβ1-4GlcNAcβ1-4GlcNAc- |
| 9 | N012 | Manα1-6(Manα1-3)Manα1-6(Neu5Acα2-3Galβ1-4GlcNAcβ1-2Manα1-3)Manβ1-4GlcNAcβ1-4GlcNAc- |
| 10 | N013 | Manα1-6(Manα1-3)Manα1-6(Neu5Acα2-6Galβ1-4GlcNAcβ1-2Manα1-3)Manβ1-4GlcNAcβ1-4GlcNAc- |
| 11 | N014 | Manα1-6(Manα1-3)Manα1-6(Galβ1-4(Fucα1-3)GlcNAcβ1-2Manα1-3)Manβ1-4GlcNAcβ1-4GlcNAc- |
| 12 | N015 | Manα1-6(Manα1-3)Manα1-6(Neu5Acα2-3Galβ1-4(Fucα1-3)GlcNAcβ1-2Manα1-3)Manβ1-4GlcNAcβ1-4GlcNAc- |
| 13 | N020 | GlcNAcβ1-2Manα1-3Manβ1-4GlcNAcβ1-4GlcNAc- |
| 14 | N021 | Galβ1-4GlcNAcβ1-2Manα1-3Manβ1-4GlcNAcβ1-4GlcNAc- |
| 15 | N022 | Neu5Acα2-3Galβ1-4GlcNAcβ1-2Manα1-3Manβ1-4GlcNAcβ1-4GlcNAc- |
| 16 | N023 | Neu5Acα2-6Galβ1-4GlcNAcβ1-2Manα1-3Manβ1-4GlcNAcβ1- |

| | | |
|---|---|---|
| | | 4GlcNAc- |
| 17 | N024 | Galβ1-4(Fucα1-3)GlcNAcβ1-2Manα1-3Manβ1-4GlcNAcβ1-4GlcNAc- |
| 18 | N025 | Neu5Acα2-3Galβ1-4(Fucα1-3)GlcNAcβ1-2Manα1-3Manβ1-4GlcNAcβ1-4GlcNAc- |
| 19 | N030 | Manα1-6(GlcNAcβ1-2Manα1-3)Manβ1-4GlcNAcβ1-4GlcNAc- |
| 20 | N031 | Manα1-6(Galβ1-4GlcNAcβ1-2Manα1-3)Manβ1-4GlcNAcβ1-4GlcNAc- |
| 21 | N032 | Manα1-6(Neu5Acα2-3Galβ1-4GlcNAcβ1-2Manα1-3)Manβ1-4GlcNAcβ1-4GlcNAc- |
| 22 | N033 | Manα1-6(Neu5Acα2-6Galβ1-4GlcNAcβ1-2Manα1-3)Manβ1-4GlcNAcβ1-4GlcNAc- |
| 23 | N034 | Manα1-6(Galβ1-4(Fucα1-3)GlcNAcβ1-2Manα1-3)Manβ1-4GlcNAcβ1-4GlcNAc- |
| 24 | N035 | Manα1-6(Neu5Acα2-3Galβ1-4(Fucα1-3)GlcNAcβ1-2Manα1-3)Manβ1-4GlcNAcβ1-4GlcNAc- |
| 25 | N040 | GlcNAcβ1-2Manα1-6Manβ1-4GlcNAcβ1-4GlcNAc- |
| 26 | N041 | Galβ1-4GlcNAcβ1-2Manα1-6Manβ1-4GlcNAcβ1-4GlcNAc- |
| 27 | N042 | Neu5Acα2-3Galβ1-4GlcNAcβ1-2Manα1-6Manβ1-4GlcNAcβ1-4GlcNAc- |
| 28 | N043 | Neu5Acα2-6Galβ1-4GlcNAcβ1-2Manα1-6Manβ1-4GlcNAcβ1-4GlcNAc- |
| 29 | N044 | Galβ1-4(Fucα1-3)GlcNAcβ1-2Manα1-6Manβ1-4GlcNAcβ1-4GlcNAc- |
| 30 | N045 | Neu5Acα2-3Galβ1-4(Fucα1-3)GlcNAcβ1-2Manα1-6Manβ1-4GlcNAcβ1-4GlcNAc- |
| 31 | N050 | GlcNAcβ1-2Manα1-6(Manα1-3)Manβ1-4GlcNAcβ1-4GlcNAc- |
| 32 | N051 | Galβ1-4GlcNAcβ1-2Manα1-6(Manα1-3)Manβ1-4GlcNAcβ1-4GlcNAc- |
| 33 | N052 | Neu5Acα2-3Galβ1-4GlcNAcβ1-2Manα1-6(Manα1-3)Manβ1-4GlcNAcβ1-4GlcNAc- |

| | | |
|---|---|---|
| 34 | N053 | Neu5Acα2-6Galβ1-4GlcNAcβ1-2Manα1-6(Manα1-3)Manβ1-4GlcNAcβ1-4GlcNAc- |
| 35 | N054 | Galβ1-4(Fucα1-3)GlcNAcβ1-2Manα1-6(Manα1-3)Manβ1-4GlcNAcβ1-4GlcNAc- |
| 36 | N055 | Neu5Acα2-3Galβ1-4(Fucα1-3)GlcNAcβ1-2Manα1-6(Manα1-3)Manβ1-4GlcNAcβ1-4GlcNAc- |
| 37 | N110 | (OAc)4GlcNAcβ1-2Manα1-6(GlcNAcβ1-2Manα1-3)Manβ1-4GlcNAcβ1-4GlcNAc- |
| 38 | N111 | GlcNAcβ1-2Manα1-6(Galβ1-4GlcNAcβ1-2Manα1-3)Manβ1-4GlcNAcβ1-4GlcNAc- |
| 39 | N112 | GlcNAcβ1-2Manα1-6(Neu5Acα2-3Galβ1-4GlcNAcβ1-2Manα1-3)Manβ1-4GlcNAcβ1-4GlcNAc- |
| 40 | N113 | GlcNAcβ1-2Manα1-6(Neu5Acα2-6Galβ1-4GlcNAcβ1-2Manα1-3)Manβ1-4GlcNAcβ1-4GlcNAc- |
| 41 | N114 | GlcNAcβ1-2Manα1-6(Galβ1-4(Fucα1-3)GlcNAcβ1-2Manα1-3)Manβ1-4GlcNAcβ1-4GlcNAc- |
| 42 | N115 | GlcNAcβ1-2Manα1-6(Neu5Acα2-3Galβ1-4(Fucα1-3)GlcNAcβ1-2Manα1-3)Manβ1-4GlcNAcβ1-4GlcNAc- |
| 43 | N122 | Galβ1-4GlcNAcβ1-2Manα1-6(Neu5Acα2-3Galβ1-4GlcNAcβ1-2Manα1-3)Manβ1-4GlcNAcβ1-4GlcNAc- |
| 44 | N123 | Galβ1-4GlcNAcβ1-2Manα1-6(Neu5Acα2-6Galβ1-4GlcNAcβ1-2Manα1-3)Manβ1-4GlcNAcβ1-4GlcNAc- |
| 45 | N124 | Galβ1-4GlcNAcβ1-2Manα1-6(Galβ1-4(Fucα1-3)GlcNAcβ1-2Manα1-3)Manβ1-4GlcNAcβ1-4GlcNAc- |
| 46 | N125 | Galβ1-4GlcNAcβ1-2Manα1-6(Neu5Acα2-3Galβ1-4(Fucα1-3)GlcNAcβ1-2Manα1-3)Manβ1-4GlcNAcβ1-4GlcNAc- |
| 47 | N133 | Neu5Acα2-3Galβ1-4GlcNAcβ1-2Manα1-6(Neu5Acα2-6Galβ1-4GlcNAcβ1-2Manα1-3)Manβ1-4GlcNAcβ1-4GlcNAc- |
| 48 | N134 | Neu5Acα2-3Galβ1-4GlcNAcβ1-2Manα1-6(Galβ1-4(Fucα1-3)GlcNAcβ1-2Manα1-3)Manβ1-4GlcNAcβ1-4GlcNAc- |
| 49 | N135 | Neu5Acα2-3Galβ1-4GlcNAcβ1-2Manα1-6(Neu5Acα2-3Galβ1-4(Fucα1-3)GlcNAcβ1-2Manα1-3)Manβ1-4GlcNAcβ1-4GlcNAc- |

| | | |
|---|---|---|
| 50 | N144 | Neu5Acα2-6Galβ1-4GlcNAcβ1-2Manα1-6(Galβ1-4(Fucα1-3)GlcNAcβ1-2Manα1-3)Manβ1-4GlcNAcβ1-4GlcNAc- |
| 51 | N155 | Galβ1-4(Fucα1-3)GlcNAcβ1-2Manα1-6(Neu5Acα2-3Galβ1-4(Fucα1-3)GlcNAcβ1-2Manα1-3)Manβ1-4GlcNAcβ1-4GlcNAc- |
| 52 | N210 | GlcNAcβ1-2Manα1-6((OAc)4GlcNAcβ1-2)Manα1-3)Manβ1-4GlcNAcβ1-4GlcNAc- |
| 53 | N211 | Galβ1-4GlcNAcβ1-2Manα1-6(GlcNAcβ1-2Manα1-3)Manβ1-4GlcNAcβ1-4GlcNAc- |
| 54 | N212 | Neu5Acα2-3Galβ1-4GlcNAcβ1-2Manα1-6(GlcNAcβ1-2Manα1-3)Manβ1-4GlcNAcβ1-4GlcNAc- |
| 55 | N213 | Neu5Acα2-6Galβ1-4GlcNAcβ1-2Manα1-6(GlcNAcβ1-2Manα1-3)Manβ1-4GlcNAcβ1-4GlcNAc- |
| 56 | N214 | Galβ1-4(Fucα1-3)GlcNAcβ1-2Manα1-6(GlcNAcβ1-2Manα1-3)Manβ1-4GlcNAcβ1-4GlcNAc- |
| 57 | N215 | Neu5Acα2-3Galβ1-4(Fucα1-3)GlcNAcβ1-2Manα1-6(GlcNAcβ1-2Manα1-3)Manβ1-4GlcNAcβ1-4GlcNAc- |
| 58 | N222 | Neu5Acα2-3Galβ1-4GlcNAcβ1-2Manα1-6(Galβ1-4GlcNAcβ1-2Manα1-3)Manβ1-4GlcNAcβ1-4GlcNAc- |
| 59 | N223 | Neu5Acα2-6Galβ1-4GlcNAcβ1-2Manα1-6(Galβ1-4GlcNAcβ1-2Manα1-3)Manβ1-4GlcNAcβ1-4GlcNAc- |
| 60 | N224 | Galβ1-4(Fucα1-3)GlcNAcβ1-2Manα1-6(Galβ1-4GlcNAcβ1-2Manα1-3)Manβ1-4GlcNAcβ1-4GlcNAc- |
| 61 | N225 | Neu5Acα2-3Galβ1-4(Fucα1-3)GlcNAcβ1-2Manα1-6(Galβ1-4GlcNAcβ1-2Manα1-3)Manβ1-4GlcNAcβ1-4GlcNAc- |
| 62 | N233 | Neu5Acα2-6Galβ1-4GlcNAcβ1-2Manα1-6(Neu5Acα2-3Galβ1-4GlcNAcβ1-2Manα1-3)Manβ1-4GlcNAcβ1-4GlcNAc- |
| 63 | N234 | Galβ1-4(Fucα1-3)GlcNAcβ1-2Manα1-6(Neu5Acα2-3Galβ1-4GlcNAcβ1-2Manα1-3)Manβ1-4GlcNAcβ1-4GlcNAc- |
| 64 | N244 | Galβ1-4(Fucα1-3)GlcNAcβ1-2Manα1-6(Neu5Acα2-6Galβ1-4GlcNAcβ1-2Manα1-3)Manβ1-4GlcNAcβ1-4GlcNAc- |
| 65 | N255 | Neu5Acα2-3Galβ1-4(Fucα1-3)GlcNAcβ1-2Manα1-6(Galβ1-4(Fucα1-3)GlcNAcβ1-2Manα1-3)Manβ1-4GlcNAcβ1-4GlcNAc- |

| | | |
|---|---|---|
| 66 | N6000 | GlcNAcβ1-2Manα1-6(GlcNAcβ1-2Manα1-3)Manβ1-4GlcNAcβ1-4(Fucα1-6)GlcNAc- |
| 67 | N6030 | Manα1-6(GlcNAcβ1-2Manα1-3)Manβ1-4GlcNAcβ1-4(Fucα1-6)GlcNAc- |
| 68 | N6111 | GlcNAcβ1-2Manα1-6(Galβ1-4GlcNAcβ1-2Manα1-3)Manβ1-4GlcNAcβ1-4(Fucα1-6)GlcNAc- |
| 69 | N6112 | GlcNAcβ1-2Manα1-6(Neu5Acα2-3Galβ1-4GlcNAcβ1-2Manα1-3)Manβ1-4GlcNAcβ1-4(Fucα1-6)GlcNAc- |
| 70 | N6113 | GlcNAcβ1-2Manα1-6(Neu5Acα2-6Galβ1-4GlcNAcβ1-2Manα1-3)Manβ1-4GlcNAcβ1-4(Fucα1-6)GlcNAc- |
| 71 | N6122 | Galβ1-4GlcNAcβ1-2Manα1-6(Neu5Acα2-3Galβ1-4GlcNAcβ1-2Manα1-3)Manβ1-4GlcNAcβ1-4(Fucα1-6)GlcNAc- |
| 72 | N6123 | Galβ1-4GlcNAcβ1-2Manα1-6(Neu5Acα2-6Galβ1-4GlcNAcβ1-2Manα1-3)Manβ1-4GlcNAcβ1-4(Fucα1-6)GlcNAc- |
| 73 | N6144 | Neu5Acα2-6Galβ1-4GlcNAcβ1-2Manα1-6(Galβ1-4(Fucα1-3)GlcNAcβ1-2Manα1-3)Manβ1-4GlcNAcβ1-4(Fucα1-6)GlcNAc- |
| 74 | N6211 | Galβ1-4GlcNAcβ1-2Manα1-6(GlcNAcβ1-2Manα1-3)Manβ1-4GlcNAcβ1-4(Fucα1-6)GlcNAc- |
| 75 | N6212 | Neu5Acα2-3Galβ1-4GlcNAcβ1-2Manα1-6(GlcNAcβ1-2Manα1-3)Manβ1-4GlcNAcβ1-4(Fucα1-6)GlcNAc- |
| 76 | N6213 | Neu5Acα2-6Galβ1-4GlcNAcβ1-2Manα1-6(GlcNAcβ1-2Manα1-3)Manβ1-4GlcNAcβ1-4(Fucα1-6)GlcNAc- |
| 77 | N6222 | Neu5Acα2-3Galβ1-4GlcNAcβ1-2Manα1-6(Galβ1-4GlcNAcβ1-2Manα1-3)Manβ1-4GlcNAcβ1-4(Fucα1-6)GlcNAc- |
| 78 | N6223 | Neu5Acα2-6Galβ1-4GlcNAcβ1-2Manα1-6(Galβ1-4GlcNAcβ1-2Manα1-3)Manβ1-4GlcNAcβ1-4(Fucα1-6)GlcNAc- |
| 79 | N6244 | Galβ1-4(Fucα1-3)GlcNAcβ1-2Manα1-6(Neu5Acα2-6Galβ1-4GlcNAcβ1-2Manα1-3)Manβ1-4GlcNAcβ1-4(Fucα1-6)GlcNAc- |
| 80 | N3001 | Galβ1-4GlcNAcβ1-2Manα1-6(Galβ1-4GlcNAcβ1-2(Galβ1-4GlcNAcβ1-4)Manα1-3)Manβ1-4GlcNAcβ1-4GlcNAc- |
| 81 | N3004 | Galβ1-4(Fucα1-3)GlcNAcβ1-2Manα1-6(Galβ1-4(Fucα1-3)GlcNAcβ1-2(Galβ1-4(Fucα1-3)GlcNAcβ1-4)Manα1-3)Manβ1-4GlcNAcβ1-4GlcNAc- |

| | | |
|---|---|---|
| 82 | Man-1 | Manβ1-4GlcNAcβ1-4GlcNAc- |
| 83 | Man-2A | Manα1-6Manβ1-4GlcNAcβ1-4GlcNAc- |
| 84 | Man-2B | Manα1-3Manβ1-4GlcNAcβ1-4GlcNAc- |
| 85 | Man-3 | Manα1-6(Manα1-3)Manβ1-4GlcNAcβ1-4GlcNAc- |
| 86 | Man-5 | Manα1-6(Manα1-3)Manα1-6(Manα1-3)Manβ1-4GlcNAcβ1-4GlcNAc- |
| 87 | Man-6 | Manα1-6(Manα1-3)Manα1-6(Manα1-2Manα1-3)Manβ1-4GlcNAcβ1-4GlcNAc- |
| 88 | Man-6D2 | Manα1-6(Manα1-2Manα1-3)Manα1-6(Manα1-3)Manβ1-4GlcNAcβ1-4GlcNAc- |
| 89 | Man-7 | [Manα1-2]Manα1-6(Manα1-3)Manα1-6(Manα1-2Manα1-3)Manβ1-4GlcNAcβ1-4GlcNAc-(3isomers) |
| 90 | Man-8 | [Manα1-2][Manα1-2]Manα1-6(Manα1-3)Manα1-6(Manα1-2Manα1-3)Manβ1-4GlcNAcβ1-4GlcNAc-(3isomers) |
| 91 | Man-9 | Manα1-2Manα1-6(Manα1-2Manα1-3)Manα1-6(Manα1-2Manα1-2Manα1-3)Manβ1-4GlcNAcβ1-4GlcNAc- |
| 92 | N002G | Neu5Gcα2-3Galβ1-4GlcNAcβ1-2Manα1-6(Neu5Gcα2-3Galβ1-4GlcNAcβ1-2Manα1-3)Manβ1-4GlcNAcβ1-4GlcNAc- |
| 93 | N003G | Neu5Gcα2-6Galβ1-4GlcNAcβ1-2Manα1-6(Neu5Gcα2-6Galβ1-4GlcNAcβ1-2Manα1-3)Manβ1-4GlcNAcβ1-4GlcNAc- |
| 94 | N012G | Manα1-6(Manα1-3)Manα1-6(Neu5Gcα2-3Galβ1-4GlcNAcβ1-2Manα1-3)Manβ1-4GlcNAcβ1-4GlcNAc- |
| 95 | N013G | Manα1-6(Manα1-3)Manα1-6(Neu5Gcα2-6Galβ1-4GlcNAcβ1-2Manα1-3)Manβ1-4GlcNAcβ1-4GlcNAc- |
| 96 | N015G | Manα1-6(Manα1-3)Manα1-6(Neu5Gcα2-3Galβ1-4(Fucα1-3)GlcNAcβ1-2Manα1-3)Manβ1-4GlcNAcβ1-4GlcNAc- |
| 97 | N022G | Neu5Gcα2-3Galβ1-4GlcNAcβ1-2Manα1-3Manβ1-4GlcNAcβ1-4GlcNAc- |
| 98 | N025G | Neu5Gcα2-3Galβ1-4(Fucα1-3)GlcNAcβ1-2Manα1-3Manβ1-4GlcNAcβ1-4GlcNAc- |
| 99 | N112G | GlcNAcβ1-2Manα1-6(Neu5Gcα2-3Galβ1-4GlcNAcβ1-2Manα1-3)Manβ1-4GlcNAcβ1-4GlcNAc- |

| | | |
|---|---|---|
| 100 | N113G | GlcNAcβ1-2Manα1-6(Neu5Gcα2-6Galβ1-4GlcNAcβ1-2Manα1-3)Manβ1-4GlcNAcβ1-4GlcNAc- |

**Table S2.** The numbering and structures of the sialylated N-glycan microarray.

| No. | Structure |
|---|---|
| GC-1 | Neu5Gcα2-3Galβ1-4GlcNAcβ1-2Manα1-6(Neu5Gcα2-3Galβ1-4GlcNAcβ1-2Manα1-3)Manβ1-4GlcNAcβ1-4GlcNAc- |
| GC-2 | Neu5Gcα2-6Galβ1-4GlcNAcβ1-2Manα1-6(Neu5Gcα2-6Galβ1-4GlcNAcβ1-2Manα1-3)Manβ1-4GlcNAcβ1-4GlcNAc- |
| GC-3 | Neu5Gcα2-3Galβ1-4(Fucα1-3)GlcNAcβ1-2Manα1-6(Neu5Gcα2-3Galβ1-4(Fucα1-3)GlcNAcβ1-2Manα1-3)Manβ1-4GlcNAcβ1-4GlcNAc- |
| GC-4 | Manα1-6(Manα1-3)Manα1-6(Neu5Gcα2-3Galβ1-4GlcNAcβ1-2Manα1-3)Manβ1-4GlcNAcβ1-4GlcNAc- |
| GC-5 | Manα1-6(Manα1-3)Manα1-6(Neu5Gcα2-6Galβ1-4GlcNAcβ1-2Manα1-3)Manβ1-4GlcNAcβ1-4GlcNAc- |
| GC-6 | Manα1-6(Manα1-3)Manα1-6(Neu5Gcα2-3Galβ1-4(Fucα1-3)GlcNAcβ1-2Manα1-3)Manβ1-4GlcNAcβ1-4GlcNAc- |
| GC-7 | Neu5Gcα2-3Galβ1-4GlcNAcβ1-2Manα1-3Manβ1-4GlcNAcβ1-4GlcNAc- |
| GC-8 | Neu5Gcα2-6Galβ1-4GlcNAcβ1-2Manα1-3Manβ1-4GlcNAcβ1-4GlcNAc- |
| GC-9 | Neu5Gcα2-3Galβ1-4(Fucα1-3)GlcNAcβ1-2Manα1-3Manβ1-4GlcNAcβ1-4GlcNAc- |
| GC-10 | Manα1-6(Neu5Gcα2-3Galβ1-4GlcNAcβ1-2Manα1-3)Manβ1-4GlcNAcβ1-4GlcNAc- |
| GC-11 | Manα1-6(Neu5Gcα2-6Galβ1-4GlcNAcβ1-2Manα1-3)Manβ1-4GlcNAcβ1-4GlcNAc- |
| GC-12 | Neu5Gcα2-3Galβ1-4GlcNAcβ1-2Manα1-6Manβ1-4GlcNAcβ1-4GlcNAc- |
| GC-13 | Neu5Gcα2-6Galβ1-4GlcNAcβ1-2Manα1-6Manβ1-4GlcNAcβ1-4GlcNAc- |
| GC-14 | Neu5Gcα2-3Galβ1-4(Fucα1-3)GlcNAcβ1-2Manα1-6Manβ1-4GlcNAcβ1-4GlcNAc- |

| | |
|---|---|
| GC-15 | Neu5Gcα2-3Galβ1-4GlcNAcβ1-2Manα1-6(Manα1-3)Manβ1-4GlcNAcβ1-4GlcNAc- |
| GC-16 | Neu5Gcα2-6Galβ1-4GlcNAcβ1-2Manα1-6(Manα1-3)Manβ1-4GlcNAcβ1-4GlcNAc- |
| GC-17 | Neu5Gcα2-3Galβ1-4(Fucα1-3)GlcNAcβ1-2Manα1-6(Manα1-3)Manβ1-4GlcNAcβ1-4GlcNAc- |
| GC-18 | GlcNAcβ1-2Manα1-6(Neu5Gcα2-3Galβ1-4GlcNAcβ1-2Manα1-3)Manβ1-4GlcNAcβ1-4GlcNAc- |
| GC-19 | GlcNAcβ1-2Manα1-6(Neu5Gcα2-6Galβ1-4GlcNAcβ1-2Manα1-3)Manβ1-4GlcNAcβ1-4GlcNAc- |
| GC-20 | GlcNAcβ1-2Manα1-6(Neu5Gcα2-3Galβ1-4(Fucα1-3)GlcNAcβ1-2Manα1-3)Manβ1-4GlcNAcβ1-4GlcNAc- |
| GC-21 | Galβ1-4GlcNAcβ1-2Manα1-6(Neu5Gcα2-3Galβ1-4GlcNAcβ1-2Manα1-3)Manβ1-4GlcNAcβ1-4GlcNAc- |
| GC-22 | Galβ1-4GlcNAcβ1-2Manα1-6(Neu5Gcα2-6Galβ1-4GlcNAcβ1-2Manα1-3)Manβ1-4GlcNAcβ1-4GlcNAc- |
| GC-23 | Galβ1-4GlcNAcβ1-2Manα1-6(Neu5Gcα2-3Galβ1-4(Fucα1-3)GlcNAcβ1-2Manα1-3)Manβ1-4GlcNAcβ1-4GlcNAc- |
| GC-24 | Neu5Gcα2-3Galβ1-4GlcNAcβ1-2Manα1-6(Neu5Gcα2-6Galβ1-4GlcNAcβ1-2Manα1-3)Manβ1-4GlcNAcβ1-4GlcNAc- |
| GC-25 | Neu5Gcα2-3Galβ1-4GlcNAcβ1-2Manα1-6(Galβ1-4(Fucα1-3)GlcNAcβ1-2Manα1-3)Manβ1-4GlcNAcβ1-4GlcNAc- |
| GC-26 | Neu5Gcα2-3Galβ1-4GlcNAcβ1-2Manα1-6(Neu5Gcα2-3Galβ1-4(Fucα1-3)GlcNAcβ1-2Manα1-3)Manβ1-4GlcNAcβ1-4GlcNAc- |
| GC-27 | Neu5Gcα2-6Galβ1-4GlcNAcβ1-2Manα1-6(Galβ1-4(Fucα1-3)GlcNAcβ1-2Manα1-3)Manβ1-4GlcNAcβ1-4GlcNAc- |
| GC-28 | Neu5Gcα2-6Galβ1-4GlcNAcβ1-2Manα1-6(Neu5Gcα2-3Galβ1-4(Fucα1-3)GlcNAcβ1-2Manα1-3)Manβ1-4GlcNAcβ1-4GlcNAc- |
| GC-29 | Galβ1-4(Fucα1-3)GlcNAcβ1-2Manα1-6(Neu5Gcα2-3Galβ1-4(Fucα1-3)GlcNAcβ1-2Manα1-3)Manβ1-4GlcNAcβ1-4GlcNAc- |

| | |
|---|---|
| GC-30 | Neu5Gcα2-3Galβ1-4GlcNAcβ1-2Manα1-6(GlcNAcβ1-2Manα1-3)Manβ1-4GlcNAcβ1-4GlcNAc- |
| GC-31 | Neu5Gcα2-6Galβ1-4GlcNAcβ1-2Manα1-6(GlcNAcβ1-2Manα1-3)Manβ1-4GlcNAcβ1-4GlcNAc- |
| GC-32 | Neu5Gcα2-3Galβ1-4(Fucα1-3)GlcNAcβ1-2Manα1-6(GlcNAcβ1-2Manα1-3)Manβ1-4GlcNAcβ1-4GlcNAc- |
| GC-33 | Neu5Gcα2-3Galβ1-4GlcNAcβ1-2Manα1-6(Galβ1-4GlcNAcβ1-2Manα1-3)Manβ1-4GlcNAcβ1-4GlcNAc- |
| GC-34 | Neu5Gcα2-6Galβ1-4GlcNAcβ1-2Manα1-6(Galβ1-4GlcNAcβ1-2Manα1-3)Manβ1-4GlcNAcβ1-4GlcNAc- |
| GC-35 | Neu5Gcα2-3Galβ1-4(Fucα1-3)GlcNAcβ1-2Manα1-6(Galβ1-4GlcNAcβ1-2Manα1-3)Manβ1-4GlcNAcβ1-4GlcNAc- |
| GC-36 | Neu5Gcα2-6Galβ1-4GlcNAcβ1-2Manα1-6(Neu5Gcα2-3Galβ1-4GlcNAcβ1-2Manα1-3)Manβ1-4GlcNAcβ1-4GlcNAc- |
| GC-37 | Neu5Gcα2-3Galβ1-4(Fucα1-3)GlcNAcβ1-2Manα1-6(Neu5Gcα2-3Galβ1-4GlcNAcβ1-2Manα1-3)Manβ1-4GlcNAcβ1-4GlcNAc- |
| GC-38 | Neu5Gcα2-3Galβ1-4(Fucα1-3)GlcNAcβ1-2Manα1-6(Neu5Gcα2-6Galβ1-4GlcNAcβ1-2Manα1-3)Manβ1-4GlcNAcβ1-4GlcNAc- |
| GC-39 | Neu5Gcα2-3Galβ1-4(Fucα1-3)GlcNAcβ1-2Manα1-6(Galβ1-4(Fucα1-3)GlcNAcβ1-2Manα1-3)Manβ1-4GlcNAcβ1-4GlcNAc- |
| GC-40 | Neu5Acα2-6Galβ1-4GlcNAcβ1-2Manα1-6(Neu5Gcα2-6Galβ1-4GlcNAcβ1-2Manα1-3)Manβ1-4GlcNAcβ1-4GlcNAc- |
| GC-41 | Neu5Gcα2-6Galβ1-4GlcNAcβ1-2Manα1-6(Neu5Acα2-6Galβ1-4GlcNAcβ1-2Manα1-3)Manβ1-4GlcNAcβ1-4GlcNAc- |
| AC-1 | Neu5Acα2-3Galβ1-4GlcNAcβ1-2Manα1-6(Neu5Acα2-3Galβ1-4GlcNAcβ1-2Manα1-3)Manβ1-4GlcNAcβ1-4GlcNAc- |
| AC-2 | Neu5Acα2-6Galβ1-4GlcNAcβ1-2Manα1-6(Neu5Acα2-6Galβ1-4GlcNAcβ1-2Manα1-3)Manβ1-4GlcNAcβ1-4GlcNAc- |
| AC-3 | Neu5Acα2-3Galβ1-4(Fucα1-3)GlcNAcβ1-2Manα1-6(Neu5Acα2-3Galβ1-4(Fucα1-3)GlcNAcβ1-2Manα1-3)Manβ1-4GlcNAcβ1-4GlcNAc- |

| | |
|---|---|
| AC-4 | Manα1-6(Manα1-3)Manα1-6(Neu5Acα2-3Galβ1-4GlcNAcβ1-2Manα1-3)Manβ1-4GlcNAcβ1-4GlcNAc- |
| AC-5 | Manα1-6(Manα1-3)Manα1-6(Neu5Acα2-6Galβ1-4GlcNAcβ1-2Manα1-3)Manβ1-4GlcNAcβ1-4GlcNAc- |
| AC-6 | Manα1-6(Manα1-3)Manα1-6(Neu5Acα2-3Galβ1-4(Fucα1-3)GlcNAcβ1-2Manα1-3)Manβ1-4GlcNAcβ1-4GlcNAc- |
| AC-7 | Neu5Acα2-3Galβ1-4GlcNAcβ1-2Manα1-3Manβ1-4GlcNAcβ1-4GlcNAc- |
| AC-8 | Neu5Acα2-6Galβ1-4GlcNAcβ1-2Manα1-3Manβ1-4GlcNAcβ1-4GlcNAc- |
| AC-9 | Neu5Acα2-3Galβ1-4(Fucα1-3)GlcNAcβ1-2Manα1-3Manβ1-4GlcNAcβ1-4GlcNAc- |
| AC-10 | Manα1-6(Neu5Acα2-3Galβ1-4GlcNAcβ1-2Manα1-3)Manβ1-4GlcNAcβ1-4GlcNAc- |
| AC-11 | Manα1-6(Neu5Acα2-6Galβ1-4GlcNAcβ1-2Manα1-3)Manβ1-4GlcNAcβ1-4GlcNAc- |
| AC-12 | Neu5Acα2-3Galβ1-4GlcNAcβ1-2Manα1-6Manβ1-4GlcNAcβ1-4GlcNAc- |
| AC-13 | Neu5Acα2-6Galβ1-4GlcNAcβ1-2Manα1-6Manβ1-4GlcNAcβ1-4GlcNAc- |
| AC-14 | Neu5Acα2-3Galβ1-4(Fucα1-3)GlcNAcβ1-2Manα1-6Manβ1-4GlcNAcβ1-4GlcNAc- |
| AC-15 | Neu5Acα2-3Galβ1-4GlcNAcβ1-2Manα1-6(Manα1-3)Manβ1-4GlcNAcβ1-4GlcNAc- |
| AC-16 | Neu5Acα2-6Galβ1-4GlcNAcβ1-2Manα1-6(Manα1-3)Manβ1-4GlcNAcβ1-4GlcNAc- |
| AC-17 | Neu5Acα2-3Galβ1-4(Fucα1-3)GlcNAcβ1-2Manα1-6(Manα1-3)Manβ1-4GlcNAcβ1-4GlcNAc- |
| AC-18 | GlcNAcβ1-2Manα1-6(Neu5Acα2-3Galβ1-4GlcNAcβ1-2Manα1-3)Manβ1-4GlcNAcβ1-4GlcNAc- |
| AC-19 | GlcNAcβ1-2Manα1-6(Neu5Acα2-6Galβ1-4GlcNAcβ1-2Manα1-3)Manβ1-4GlcNAcβ1-4GlcNAc- |
| AC-20 | GlcNAcβ1-2Manα1-6(Neu5Acα2-3Galβ1-4(Fucα1-3)GlcNAcβ1-2Manα1- |

| | |
|---|---|
| | 3)Manβ1-4GlcNAcβ1-4GlcNAc- |
| AC-21 | Galβ1-4GlcNAcβ1-2Manα1-6(Neu5Acα2-3Galβ1-4GlcNAcβ1-2Manα1-3)Manβ1-4GlcNAcβ1-4GlcNAc- |
| AC-22 | Galβ1-4GlcNAcβ1-2Manα1-6(Neu5Acα2-6Galβ1-4GlcNAcβ1-2Manα1-3)Manβ1-4GlcNAcβ1-4GlcNAc- |
| AC-23 | Galβ1-4GlcNAcβ1-2Manα1-6(Neu5Acα2-3Galβ1-4(Fucα1-3)GlcNAcβ1-2Manα1-3)Manβ1-4GlcNAcβ1-4GlcNAc- |
| AC-24 | Neu5Acα2-3Galβ1-4GlcNAcβ1-2Manα1-6(Neu5Acα2-6Galβ1-4GlcNAcβ1-2Manα1-3)Manβ1-4GlcNAcβ1-4GlcNAc- |
| AC-25 | Neu5Acα2-3Galβ1-4GlcNAcβ1-2Manα1-6(Galβ1-4(Fucα1-3)GlcNAcβ1-2Manα1-3)Manβ1-4GlcNAcβ1-4GlcNAc- |
| AC-26 | Neu5Acα2-3Galβ1-4GlcNAcβ1-2Manα1-6(Neu5Acα2-3Galβ1-4(Fucα1-3)GlcNAcβ1-2Manα1-3)Manβ1-4GlcNAcβ1-4GlcNAc- |
| AC-27 | Neu5Acα2-6Galβ1-4GlcNAcβ1-2Manα1-6(Galβ1-4(Fucα1-3)GlcNAcβ1-2Manα1-3)Manβ1-4GlcNAcβ1-4GlcNAc- |
| AC-29 | Galβ1-4(Fucα1-3)GlcNAcβ1-2Manα1-6(Neu5Acα2-3Galβ1-4(Fucα1-3)GlcNAcβ1-2Manα1-3)Manβ1-4GlcNAcβ1-4GlcNAc- |
| AC-30 | Neu5Acα2-3Galβ1-4GlcNAcβ1-2Manα1-6(GlcNAcβ1-2Manα1-3)Manβ1-4GlcNAcβ1-4GlcNAc- |
| AC-31 | Neu5Acα2-6Galβ1-4GlcNAcβ1-2Manα1-6(GlcNAcβ1-2Manα1-3)Manβ1-4GlcNAcβ1-4GlcNAc- |
| AC-32 | Neu5Acα2-3Galβ1-4(Fucα1-3)GlcNAcβ1-2Manα1-6(GlcNAcβ1-2Manα1-3)Manβ1-4GlcNAcβ1-4GlcNAc- |
| AC-33 | Neu5Acα2-3Galβ1-4GlcNAcβ1-2Manα1-6(Galβ1-4GlcNAcβ1-2Manα1-3)Manβ1-4GlcNAcβ1-4GlcNAc- |
| AC-34 | Neu5Acα2-6Galβ1-4GlcNAcβ1-2Manα1-6(Galβ1-4GlcNAcβ1-2Manα1-3)Manβ1-4GlcNAcβ1-4GlcNAc- |
| AC-35 | Neu5Acα2-3Galβ1-4(Fucα1-3)GlcNAcβ1-2Manα1-6(Galβ1-4GlcNAcβ1-2Manα1-3)Manβ1-4GlcNAcβ1-4GlcNAc- |
| AC-36 | Neu5Acα2-6Galβ1-4GlcNAcβ1-2Manα1-6(Neu5Acα2-3Galβ1- |

| | |
|---|---|
| | 4GlcNAcβ1-2Manα1-3)Manβ1-4GlcNAcβ1-4GlcNAc- |
| AC-39 | Neu5Acα2-3Galβ1-4(Fucα1-3)GlcNAcβ1-2Manα1-6(Galβ1-4(Fucα1-3)GlcNAcβ1-2Manα1-3)Manβ1-4GlcNAcβ1-4GlcNAc- |

**Table S3.** The numbering and structures of the O-glycan microarray.

| ID | Structure |
|---|---|
| O1 | GalNAcα-Ser |
| O2 | GalNAcα-Thr |
| O3 | Neu5Acα2-6GalNAcα-Ser |
| O4 | Neu5Acα2-6GalNAcα-Thr |
| O5 | Galβ1-3GalNAcα-Ser |
| O6 | Galβ1-3GalNAcα-Thr |
| O7 | Neu5Acα2-3Galβ1-3GalNAcα-Ser |
| O8 | Neu5Gcα2-3Galβ1-3GalNAcα-Ser |
| O9 | GalNAcβ1-4(Neu5Acα2-3)GalNAcα-Ser |
| O10 | Fucα1-2Galβ1-3GalNAcα-Ser |
| O11 | GalNAcβ1-3(Fucα1-2)Galβ1-3GalNAcα-Ser |
| O12 | Galα1-3(Fucα1-2)Galβ1-3GalNAcα-Ser |
| O13 | GlcNAcβ1-3Galβ1-3GalNAcα-Ser |
| O14 | Galβ1-4GlcNAcβ1-3Galβ1-3GalNAcα-Ser |
| O15 | Galα1-3Galβ1-4GlcNAcβ1-3Galβ1-3GalNAcα-Ser |
| O16 | Neu5Acα2-3Galβ1-4GlcNAcβ1-3Galβ1-3GalNAcα-Ser |
| O17 | GalNAcβ1-4(Neu5Acα2-3)Galβ1-4GlcNAcβ1-3Galβ1-3GalNAcα-Ser |
| O18 | Fucα1-2Galβ1-4GlcNAcβ1-3Galβ1-3GalNAcα-Ser |
| O19 | Galβ1-4(Fucα1-2)Galβ1-4GlcNAcβ1-3Galβ1-3GalNAcα-Ser |
| O20 | Fucα1-2Galβ1-4(Fucα1-2)Galβ1-4GlcNAcβ1-3Galβ1-3GalNAcα-Ser |
| O21 | Neu5Acα2-6(Neu5Acα2-3(GalNAcβ1-4)Galβ1-3)GalNAcα-Ser |

| | |
|---|---|
| O22 | GlcNAcβ1-6(Galβ1-3)GalNAcα-Ser |
| O23 | GlcNAcβ1-6(Galβ1-3)GalNAcα-Thr |
| O24 | Galβ1-4GlcNAcβ1-6(Galβ1-3)GalNAcα-Ser |
| O25 | GlcNAcβ1-3GalNAcα-Ser |
| O26 | GlcNAcβ1-3GalNAcα-Thr |
| O27 | Galβ1-4GlcNAcβ1-3GalNAcα-Ser |
| O28 | Galα1-3Galβ1-4GlcNAcβ1-3GalNAcα-Ser |
| O29 | Neu5Acα2-3Galβ1-4GlcNAcβ1-3GalNAcα-Ser |
| O30 | Neu5Acα2-6Galβ1-4GlcNAcβ1-3GalNAcα-Ser |
| O31 | GalNAcβ1-4(Neu5Acα2-3)Galβ1-4GlcNAcβ1-3GalNAcα-Ser |
| O32 | Galβ1-4(Fucα1-3)GlcNAcβ1-3GalNAcα-Ser |
| O33 | Fucα1-2Galβ1-4(Fucα1-3)GlcNAcβ1-3GalNAcα-Ser |
| O34 | Fucα1-2Galβ1-4GlcNAcβ1-3GalNAcα-Ser |
| O35 | GalNAcα1-3(Fucα1-2)GlcNAcβ1-3GalNAcα-Ser |
| O36 | Galα1-3(Fucα1-2)GlcNAcβ1-3GalNAcα-Ser |
| O37 | Neu5Acα2-6(GlcNAcβ1-3)GalNAcα-Ser |
| O38 | Neu5Acα2-6(Galβ1-4GlcNAcβ1-3)GalNAcα-Ser |
| O39 | GlcNAcβ1-6(GlcNAcβ1-3)GalNAcα-Thr |
| O40 | GlcNAcβ1-6GalNAcα-Ser |
| O41 | Galβ1-4GlcNAcβ1-6GalNAcα-Ser |
| O42 | Galα1-3Galβ1-4GlcNAcβ1-6GalNAcα-Ser |
| O43 | GalNAcβ1-4(Neu5Acα2-3)Galβ1-4GlcNAcβ1-6GalNAcα-Ser |
| O44 | Galβ1-4(Fucα1-3)GlcNAcβ1-6GalNAcα-Ser |
| O45 | Neu5Acα2-3Galβ1-4(Fucα1-3)GlcNAcβ1-6GalNAcα-Ser |
| O46 | Fucα1-2Galβ1-4(Fucα1-3)GlcNAcβ1-6GalNAcα-Ser |
| O47 | Fucα1-2Galβ1-4GlcNAcβ1-6GalNAcα-Ser |
| O48 | GalNAcα1-3(Fucα1-2)Galβ1-4GlcNAcβ1-6GalNAcα-Ser |

| | |
|---|---|
| O49 | GalNAcα-H2N-APGSTAPP-NH2 |
| O50 | GalNAcα-H2N-TSAPDTRPAP-NH2 |
| O51 | GlcNAcβ1-2Manα-Thr |
| O52 | Galβ1-4GlcNAcβ1-2Manα-Thr |
| O53 | Neu5Acα2-3Galβ1-4GlcNAcβ1-2Manα-Thr |
| O54 | Neu5Gcα2-3Galβ1-4GlcNAcβ1-2Manα-Thr |
| O55 | Neu5Acα2-6Galβ1-4GlcNAcβ1-2Manα-Thr |
| O56 | Neu5Gcα2-6Galβ1-4GlcNAcβ1-2Manα-Thr |
| O57 | Galβ1-4(Fucα1-3)GlcNAcβ1-2Manα-Thr |
| O58 | Neu5Acα2-3Galβ1-4(Fucα1-3)GlcNAcβ1-2Manα-Thr |
| O59 | GlcAβ1-3Galβ1-4GlcNAcβ1-2Manα-Thr |
| O60 | GlcNAcβ1-6(GlcNAcβ1-2)Manα-Thr |
| O61 | GlcNAcβ1-6(Galβ1-4GlcNAcβ1-2)Manα-Thr |
| O62 | GlcNAcβ1-6(Neu5Acα2-3Galβ1-4GlcNAcβ1-2)Manα-Thr |
| O63 | GlcNAcβ1-6(Neu5Acα2-6Galβ1-4GlcNAcβ1-2)Manα-Thr |
| O64 | GlcNAcβ1-6(Galβ1-4(Fucα1-3)GlcNAcβ1-2)Manα-Thr |
| O65 | GlcNAcβ1-6(Neu5Acα2-6Galβ1-4(Fucα1-3)GlcNAcβ1-2)Manα-Thr |
| O66 | Galβ1-4GlcNAcβ1-6(Galβ1-4GlcNAcβ1-2)Manα-Thr |
| O67 | Galβ1-4GlcNAcβ1-6(Neu5Acα2-3Galβ1-4GlcNAcβ1-2)Manα-Thr |
| O68 | Galβ1-4GlcNAcβ1-6(Neu5Acα2-6Galβ1-4GlcNAcβ1-2)Manα-Thr |
| O69 | Galβ1-4GlcNAcβ1-6(Galβ1-4(Fucα1-3)GlcNAcβ1-2)Manα-Thr |
| O70 | Galβ1-4GlcNAcβ1-6(Neu5Acα2-3Galβ1-4(Fucα1-3)GlcNAcβ1-2)Manα-Thr |
| O71 | Neu5Acα2-3Galβ1-4GlcNAcβ1-6(Neu5Acα2-3Galβ1-4GlcNAcβ1-2)Manα-Thr |
| O72 | Neu5Acα2-3Galβ1-4GlcNAcβ1-6(Neu5Acα2-6Galβ1-4GlcNAcβ1-2)Manα-Thr |

| | |
|---|---|
| O73 | Neu5Acα2-3Galβ1-4GlcNAcβ1-6(Galβ1-4(Fucα1-3)GlcNAcβ1-2)Manα-Thr |
| O74 | Neu5Acα2-3Galβ1-4GlcNAcβ1-6(Neu5Acα2-3Galβ1-4(Fucα1-3)GlcNAcβ1-2)Manα-Thr |
| O75 | Neu5Acα2-6Galβ1-4GlcNAcβ1-6(Neu5Acα2-6Galβ1-4GlcNAcβ1-2)Manα-Thr |
| O76 | Neu5Acα2-6Galβ1-4GlcNAcβ1-6(Galβ1-4(Fucα1-3)GlcNAcβ1-2)Manα-Thr |
| O77 | Neu5Acα2-6Galβ1-4GlcNAcβ1-6(Neu5Acα2-3Galβ1-4(Fucα1-3)GlcNAcβ1-2)Manα-Thr |
| O78 | Galβ1-4(Fucα1-3)GlcNAcβ1-6(Galβ1-4(Fucα1-3)GlcNAcβ1-2)Manα-Thr |
| O79 | Galβ1-4(Fucα1-3)GlcNAcβ1-6(Neu5Acα2-3Galβ1-4(Fucα1-3)GlcNAcβ1-2)Manα-Thr |
| O80 | Neu5Acα2-3Galβ1-4(Fucα1-3)GlcNAcβ1-6(Neu5Acα2-3Galβ1-4(Fucα1-3)GlcNAcβ1-2)Manα-Thr |
| O81 | Galβ1-4GlcNAcβ1-6(GlcNAcβ1-2)Manα-Thr |
| O82 | Neu5Acα2-6Galβ1-4GlcNAcβ1-6(GlcNAcβ1-2)Manα-Thr |
| O83 | Galβ1-4(Fucα1-3)GlcNAcβ1-6(GlcNAcβ1-2)Manα-Thr |
| O84 | Neu5Acα2-3Galβ1-4(Fucα1-3)GlcNAcβ1-6(GlcNAcβ1-2)Manα-Thr |
| O85 | Neu5Acα2-3Galβ1-4GlcNAcβ1-6(Galβ1-4GlcNAcβ1-2)Manα-Thr |
| O86 | Neu5Acα2-6Galβ1-4GlcNAcβ1-6(Galβ1-4GlcNAcβ1-2)Manα-Thr |
| O87 | Galβ1-4(Fucα1-3)GlcNAcβ1-6(Galβ1-4)GlcNAcβ1-2)Manα-Thr |
| O88 | Neu5Acα2-3Galβ1-4(Fucα1-3)GlcNAcβ1-6(Galβ1-4GlcNAcβ1-2)Manα-Thr |
| O89 | Neu5Acα2-6Galβ1-4GlcNAcβ1-6(Neu5Acα2-3Galβ1-4GlcNAcβ1-2)Manα-Thr |
| O90 | Galβ1-4(Fucα1-3)GlcNAcβ1-6(Neu5Acα2-3Galβ1-4GlcNAcβ1-2)Manα-Thr |
| O91 | Neu5Acα2-3Galβ1-4(Fucα1-3)GlcNAcβ1-6(Neu5Acα2-3Galβ1-4GlcNAcβ1-2)Manα-Thr |

| | |
|---|---|
| O92 | Galβ1-4(Fucα1-3)GlcNAcβ1-6(Neu5Acα2-6Galβ1-4GlcNAcβ1-2)Manα-Thr |
| O93 | Neu5Acα2-3Galβ1-4(Fucα1-3)GlcNAcβ1-6(Neu5Acα2-6Galβ1-4GlcNAcβ1-2)Manα-Thr |
| O94 | Neu5Acα2-3Galβ1-4(Fucα1-3)GlcNAcβ1-6(Galβ1-4(Fucα1-3)GlcNAcβ1-2)Manα-Thr |

**Table S4.** The numbering and structures of the glycolipid glycan microarray.

| ID | Glycan Structure |
|---|---|
| G1 | Neu5Acα2–3Galβ1–4Glc |
| G2 | Neu5Gcα2–3Galβ1–4Glc |
| G3 | Kdnα2–3Galβ1–4Glc |
| G4 | Neu5Ac8Meα2–3Galβ1–4Glc |
| G5 | Neu5Acα2–3(GalNAcβ1–4)Galβ1–4Glc |
| G6 | Neu5Gcα2–3(GalNAcβ1–4)Galβ1–4Glc |
| G7 | Kdnα2–3(GalNAcβ1–4)Galβ1–4Glc |
| G8 | Neu5Acα2–3(Galβ1–3GalNAcβ1–4)Galβ1–4Glc |
| G9 | Neu5Gcα2–3(Galβ1–3GalNAcβ1–4)Galβ1–4Glc |
| G10 | Kdnα2–3(Galβ1–3GalNAcβ1–4)Galβ1–4Glc |
| G11 | Neu5Acα2–8Neu5Acα2–3Galβ1–4Glc |
| G12 | Neu5Acα2–8Neu5Gcα2–3Galβ1–4Glc |
| G13 | Neu5Acα2–8Kdncα2–3Galβ1–4Glc |
| G14 | Neu5Gcα2–8Neu5Acα2–3Galβ1–4Glc |
| G15 | Neu5Gcα2–8Neu5Gcα2–3Galβ1–4Glc |
| G16 | Kdnα2–8Neu5Acα2–3Galβ1–4Glc |
| G17 | Kdnα2–8Neu5Gcα2–3Galβ1–4Glc |
| G18 | Kdnα2–8Kdnα2–3Galβ1–4Glc |
| G19 | Neu5Acα2–8Neu5Acα2–3(GalNAcβ1–4)Galβ1–4Glc |

| | |
|---|---|
| G20 | Neu5Acα2–8Neu5Gcα2–3(GalNAcβ1–4)Galβ1–4Glc |
| G21 | Neu5Gcα2–8Neu5Acα2–3(GalNAcβ1–4)Galβ1–4Glc |
| G22 | Neu5Gcα2–8Neu5Gcα2–3(GalNAcβ1–4)Galβ1–4Glc |
| G23 | Kdnα2–8Neu5Acα2–3(GalNAcβ1–4)Galβ1–4Glc |
| G24 | Kdnα2–8Neu5Gcα2–3(GalNAcβ1–4)Galβ1–4Glc |
| G25 | Kdnα2–8Kdnα2–3(GalNAcβ1–4)Galβ1–4Glc |
| G26 | Neu5Acα2–3Galβ1–3GalNAcb1–4(Neu5Aca2–3)Galβ1–4Glc |
| G27 | Neu5Acα2–8Neu5Acα2–3(Galβ1–3GalNAcβ1–4)Galβ1–4Glc |
| G28 | Neu5Gcα2–8Neu5Gcα2–3(Galβ1–3GalNAcβ1–4)Galβ1–4Glc |
| G29 | Kdnα2–8Neu5Gcα2–3(Galβ1–3GalNAcβ1–4)Galβ1–4Glc |
| G30 | Neu5Acα2–8Neu5Acα2–3Galβ1–3GalNAcβ1–4(Neu5Aca2–3)Galβ1–4Glc |
| G31 | GalNAcβ1-4(Neu5Acα2–8Neu5Acα2–8Neu5Acα2–3)Galβ1–4Glc |
| G32 | Neu5Acα2–8Neu5Acα2–8Neu5Acα2–3Galβ1–4Glc |
| G33 | GlcNAcβ1–3Galβ1–4Glc |
| G34 | Galβ1–3GlcNAcβ1–3Galβ1–4Glc |
| G35 | Galβ1–4GlcNAcβ1–3Galβ1–4Glc |
| G36 | Galβ1–4(Fucα1–3)GlcNAcβ1–3Galβ1–4Glc |
| G37 | Neu5Acα2–3Galβ1–4GlcNAcβ1–3Galβ1–4Glc |
| G38 | Neu5Gcα2–3Galβ1–4GlcNAcβ1–3Galβ1–4Glc |
| G39 | Kdnα2–3Galβ1–4GlcNAcβ1–3Galβ1–4Glc |
| G40 | Neu5Ac8Meα2–3Galβ1–4GlcNAcβ1–3Galβ1–4Glc |
| G41 | Neu5Acα2–3Galβ1–3GlcNAcβ1–3Galβ1–4Glc |
| G42 | Neu5Gcα2–3Galβ1–3GlcNAcβ1–3Galβ1–4Glc |
| G43 | Kdnα2–3Galβ1–3GlcNAcβ1–3Galβ1–4Glc |
| G44 | Neu5Ac8Meα2–3Galβ1–3GlcNAcβ1–3Galβ1–4Glc |
| G45 | Neu5Gcα2–3Galβ1–4(Fucα1–3)GlcNAcβ1–3Galβ1–4Glc |
| G46 | Kdnα2–3Galβ1–4(Fucα1–3)GlcNAcβ1–3Galβ1–4Glc |

| | |
|---|---|
| G47 | Galα1–4Galβ1–4Glc |
| G48 | Galα1–3Galβ1–4Glc |
| G49 | GalNAcβ1–3Galα1–4Galβ1–4Glc |
| G50 | GalNAcβ1–3Galα1–3Galβ1–4Glc |
| G51 | Galβ1–3GalNAcβ1–3Galα1–4Galβ1–4Glc |
| G52 | Galβ1–3GalNAcβ1–3Galα1–3Galβ1–4Glc |
| G53 | Fucα1-2Galβ1-3GalNAcβ1-3Galα1-4Galβ1-4Glc |
| G54 | Neu5Gcα2–3Galβ1–3GalNAcβ1–3Galα1–4Galβ1–4Glc |
| G55 | Kdnα2–3Galβ1–3GalNAcβ1–3Galα1–4Galβ1–4Glc |
| G56 | Neu5Acα2–3Galβ1–3GalNAcβ1–3Galα1–3Galβ1–4Glc |
| G57 | Neu5Gcα2–3Galβ1–3GalNAcβ1–3Galα1–3Galβ1–4Glc |
| G58 | Kdnα2–3Galβ1–3GalNAcβ1–3Galα1–3Galβ1–4Glc |

**Table S5.** Comparison of the binding affinities determined by three research teams.

| Protein | Binding affinities between heparin and CoV S proteins | | |
|---|---|---|---|
| | **Linhardt** | **Boons** | **Tan** |
| SARS-CoV2-RBD | | 1.0 μM | 239.9 μM |
| SARS-CoV-2-S1 | | | 43.3 μM |
| SARS-CoV-2-S2 | | | 6.9 μM |
| SARS-CoV-2-S | 40 pM | 55 nM | 2.2 μM |
| SARS-CoV-2-S-trimer | 73 pM | | 16.1μM |
| SARS-CoV-RBD | | | 30.5 μM |
| SARS-CoV-S1 | | | 9.9 μM |
| SARS-CoV-S | 500 nM | | 6.7 μM |
| MERS-CoV-RBD | | | 98.9 μM |
| MERS-CoV-S1 | | | 4.8 μM |
| MERS-CoV-S2 | | | 3.4 μM |
| MERS-CoV-S | 1 nM | | 21.5 μM |

**Binding of recombinant proteins to N-, O-, and glycolipid glycan microarrays.** The tested protein molecules were incubated at different concentrations with microarrays. SARS-CoV-2-RBD were incubated with the N-glycan microarray at concentrations of 1.25 μg/ml, 2.5 μg/ml, 5 μg/ml and 10 μg/ml; SARS-CoV-2-S1 were incubated with the N-glycan microarray at concentrations of 0.5 μg/ml, 1 μg/ml, 2 μg/ml and 4 μg/ml; SARS-CoV-2-S were incubated with the sialylated N-, O-, and glycolipid glycan microarrays at concentrations of 2 μg/ml and 4 μg/ml; SARS-CoV-S were incubated with the sialylated N-, O-, and glycolipid glycan microarrays at concentrations of 2 μg/ml and 4 μg/ml; MERS-CoV-S were incubated with the sialylated N-, O-, and glycolipid glycan microarrays at concentrations of 2 μg/ml and 4 μg/ml.
